# Negative Cooperativity between Gemin2 and RNA provides Insights into RNA Selection and the SMN Complex’s Release in snRNP Assembly

**DOI:** 10.1101/312124

**Authors:** Hongfei Yi, Li Mu, Congcong Shen, Xi Kong, Yingzhi Wang, Yan Hou, Rundong Zhang

## Abstract

The assembly of snRNP cores, in which seven Sm proteins, D1/D2/F/E/G/D3/B, form a ring around the nonameric Sm site of snRNAs, is the early step of spliceosome formation and essential to eukaryotes. It is mediated by the PMRT5 and SMN complexes sequentially *in vivo*. SMN deficiency causes neurodegenerative disease spinal muscular atrophy (SMA). How the SMN complex assembles snRNP cores is largely unknown, especially how the SMN complex achieves high RNA assembly specificity and how it is released. Here we show, using crystallographic and biochemical approaches, that Gemin2 of the SMN complex enhances RNA specificity of SmD1/D2/F/E/G via a negative cooperativity between Gemin2 and RNA in binding SmD1/D2/F/E/G. Gemin2, independent of its N-tail, constrains the horseshoe-shaped SmD1/D2/F/E/G from outside in a physiologically relevant, narrow state, enabling high RNA specificity. Moreover, the assembly of RNAs inside widens SmD1/D2/F/E/G, causes the release of Gemin2/SMN allosterically and allows SmD3/B to join. The assembly of SmD3/B further facilitates the release of Gemin2/SMN. This is the first to show negative cooperativity in snRNP assembly, which provides insights into RNA selection and the SMN complex’s release. These findings reveal a basic mechanism of snRNP core assembly and facilitate pathogenesis studies of SMA.

## INTRODUCTION

Small nuclear ribonucleoprotein particles (snRNPs) are major building blocks of the spliceosome, which carries out precursor mRNA splicing in eukaryotes. All snRNPs share a common feature: seven Sm (D1, D2, F, E, G, D3 and B) or Sm-like proteins (Lsm2-8) form a ring around a segment of the small nuclear RNA (snRNA) after which the snRNP is named. Correspondingly, the snRNPs can be divided into two classes: Sm-class snRNPs (U1, U2, U4 and U5 snRNPs for the major spliceosome, and U11, U12, U4atac and U5 for the minor spliceosome) and Sm-like-class snRNPs (U6 and U6atac snRNPs) (1, 2). In addition, their assemblies also take different pathways. While Sm-like-class snRNPs are assembled completely inside the nucleus and without the assistance of assembly chaperones, Sm-class snRNPs are assembled in both the nucleus and cytoplasm, and are mediated by a number of assembly chaperones (2, 3). After being transcribed in the nucleus, precursor snRNAs (pre-snRNAs) are exported into the cytoplasm, where seven Sm proteins are assembled on the Sm site, PuAUUUNUGPu, of the RNAs to form snRNP cores (Sm cores). Proper assembly of the Sm core is pre-required for hypermethylation of snRNA’s cap and import into the nucleus. After import into the nucleus, Sm-class snRNPs are maturated by further modification of RNA and joining of proteins specific to individual snRNP before they participate in pre-mRNA splicing.

Sm core assembly is a pivotal step of snRNP biogenesis and essential for eukaryotes (4). Early studies established that Sm core assembly can occur spontaneously *in vitro* by mixing the three Sm hetero-oligomers, SmD1/D2, SmF/E/G and SmD3/B, with snRNA, or even oligoribonucleotide containing just the nonameric Sm site (5, 6). The reaction takes a stepwise fashion. SmD1/D2 and SmF/E/G bind RNA to form a stable subcore, and then SmD3/B joins to form a highly stable Sm core (5). Interestingly, inside cells, Sm core assembly is mediated by a number of assembly chaperone proteins, classified into two complexes, the PRMT5 (protein arginine methyltransferase 5 complex, including 3 proteins: PRMT5, WD45 and pICln) and SMN complexes (survival motor neuron complex, including 9 proteins: SMN, Gemin2-8, and unrip) in vertebrates (2, 7). Since cells contain many RNAs which would have sequences resembling the nonameric Sm site, Sm core can potentially assemble on many illicit RNAs and cause deleterious consequence (8). The assembly chaperones, especially the SMN complex (8), are found to confer highly specific Sm core assembly, ensuring Sm proteins to assemble exclusively on cognate snRNAs, which contain both the nonameric Sm site and a 3’-adjacent stem-loop (SL), altogether termed as the snRNP code (9).

The two complexes perform assembly chaperoning roles in consecutive phases. In the first phase, PRMT5/WD45 methylate the C-terminal arginine residues of SmD3, SmB and SmD1, which is believed to enhance the interactions between Sm proteins and SMN (10, 11). pICln recruits SmD1/D2 and SmF/E/G to form a ring-shaped 6S complex, which pre-arranges the 5 Sm proteins in the finally assembled order and simultaneously prevents the entry of any RNAs to the RNA-binding pocket (12, 13). In addition, pICln also binds SmD3/B (12, 13). In the second phase, the SMN complex accepts SmD1/D2/F/E/G (5Sm) and SmD3/B and releases pICln (12). Gemin2 is the acceptor of 5Sm (14, 15). SMN binds Gemin2 by its N-terminal Gemin2-binding domain (Ge2BD, residues 26-62) (14). Both SMN and Gemin2 are highly conserved in eukaryotes (16). Either *smn* or *Gemin2* gene knockout causes early embryonic death in vertebrates, indicating the essential roles of the SMN complex in eukaryotic cells (17, 18). Moreover, the deficiency of SMN causes human neurodegenerative disease spinal muscular atrophy (SMA) (19–21), and knockdown of Gemin2 causes motor neuron degeneration in zebrafish (22) and depresses motoric abilities in Drosophila (23), emphasizing the pathophysiological relevance of the Sm-core assembly pathway. Therefore, understanding the mechanism of Sm core assembly, especially at the second phase, is of great importance because of both its fundamental role in gene expression and its potential application in SMA therapy. SMN also interacts with Gemin8 by its C-terminal self-oligomized YG box (24) and Gemin8 further binds Gemin6/7 and Unrip, but their roles are poorly understood (24–26). Gemin3 contains a DEAD box domain and is a putative RNA helicase (27). Gemin4 usually forms a complex with Gemin3, but its role is unknown (28). Gemin5 is the component to initially bind pre-snRNAs and deliver them to the rest of the SMN complex for assembly into the Sm core (29), and is currently considered to be the protein conferring the RNA assembly specificity by direct recognition of the snRNP code (30–35).

Recent structural studies of some assembled and intermediate complexes of Sm core assembly have provided great insights into the mechanisms of this complicated process. The assembled structures of U1 snRNPs and U4 snRNP cores explain how the nonameric Sm site RNA interacts specifically with the seven Sm proteins (36–40). The structures of the 6S complex (human 5Sm plus Drosophila pICln), the 8S complex (6S plus Drosophila Gemin2/SMN-N-terminal domain) and the later phase human SMN(26–62)/Gemin2/5Sm complex (hereafter we will refer to it as the 7S complex for brevity because it is equivalent to the 7S complex reported earlier which contains additional segments of SMN(12)) provide detailed insights into the mechanisms of the first phase, the transition from the first phase to the second phase, and the initial state of the second phase(14, 15). Just recently, the structures of the complexes comprised of Gemin5’s N-terminal WD domain and RNA provide details of the recognition of RNA by Gemin5 (32–34).

Despite these advances in understanding the mechanisms of Sm core assembly, there are still several important questions unanswered or not well explained, especially in the second phase mediated by the SMN complex. Question one is, how does the SMN complex enhance RNA assembly specificity? This is the central question of chaperone-mediated Sm core assembly because Sm cores can form *in vitro* spontaneously and specifically on RNAs containing just the nonameric Sm site and it is the SMN complex that enhances the specificity of Sm core assembly (8, 9). Although current knowledge considers that Gemin5 is the right protein by direct binding to the snRNP code and this model is partially supported by some experimental data (30–35), there are several conflicting observations indicating that the problem is not solved yet. First, Sm core assembly is a highly conserved pathway in all eukaryotes, but there is no homolog of Gemin5 in many lower eukaryotes (16, 35). Second, recent structural and biochemical studies showed that the RNA-binding specificity of Gemin5 is only able to recognize part of the Sm site, AUUU, not to mention the full feature of the snRNP code (32–34). Third, Gemin5 can bind promiscuous RNAs, i.e., U1-tfs which are truncated U1 pre-snRNAs lacking the Sm site and the following SL (32). Question two is, how is the SMN complex released from the Sm core? In the spliceosome, the mature snRNPs do not contain any component of the SMN complex (41, 42), but most proteins of this complex have been observed to enter the nucleus and concentrate on Cajal bodies (CBs) although in small percentage (29). Moreover, our previous 7S complex structure and biochemical tests show that Gemin2 tightly binds to 5Sm (14). How the SMN complex dissociates from the mature Sm cores is unknown.

In this study, we examined the assembly reactions in the second phase mediated by the SMN complex, from the initial state of the 7S complex formation to the completion of the Sm core, by a combination of crystallographic and biochemical approaches. We found that Gemin2 and RNA bind 5Sm in a negative cooperative manner. Independent of its N-terminal segment (residue 1-39), Gemin2, binding outside 5Sm, constrains 5Sm in a narrow state, which enhances RNA specificity, selecting the cognate snRNAs, containing the snRNP code, to assemble into the Sm subcore. snRNAs’ assembly inside 5Sm widens 5Sm, unexpectedly causing Gemin2’s release. 5Sm’s widening also allows SmD3/B to join to complete the Sm core, which further expedites Gemin2’s release. Although the concept of negative cooperativity is well known, it has never been described in snRNP assembly before. These results reveal negative cooperativity in snRNP core assembly for the first time, provide deep insights into the mechanism of Sm core assembly, mainly on RNA selection and the release of the SMN complex, and facilitate pathogenesis studies of SMA. In addition, the biogenesis of U7 snRNP, which is required for the 3’-end processing of histone mRNAs in metazoans, follows the same maturation pathway as the spliceosomal Sm-class snRNPs. The assembly of its core is also mediated by the SMN complex and requires a 3’-SL adjacent to the Sm site, although U7 snRNP core has Lsm10/11 heterodimer instead of SmD1/D2 and a slightly different Sm site, AAUUUGU<u>CUAG</u> (the major difference underlined), in U7 snRNA(43–45). We predict that the negative cooperativity mechanism likewise apply to this process.

## MATERIALS AND METHODS

### Plasmid Construction and Protein Expression and Purification

All of the plasmids used in the studies contain human complementary DNAs (cDNAs). Full-length SmD1 and SmD2 (pCDFDuet-HT-D2-D1), full-length SmF and SmE (pCDFDuet-HT-F-E), full-length SmG (pET28-HT-G) and full-length Gemin2 (pCDF-HT-Gemin2) were constructed as described before (14). SmD1s (residues 1-82) and SmD2 (pCDFDuet-HT-D2-D1s) were constructed by replacing the full-length D1 with SmD1s in pCDFDuet-HT-D2-D1. The Sm fold portion of SmD3(residues 1-75) and SmB(residues 1-91) [pCDFDuet-HT-B(1-91)-D3(1-75)] were constructed in a single pCDFDuet vector (Novagen) with N-terminal His(6)-tag followed by Tobacco Etch Virus (TEV) cleavage site (HT) fused to SmB. Gemin2ΔN39 (pCDF-HT-Gemin2ΔN39) were constructed by deletion of the N-terminal 39 residues in pCDF-HT-Gemin2. SMN_Ge2BD_, containing SMN residues 26–62 (pET21-HMT-SMN_Ge2BD_), was fused with an N-terminal His(6)-tag followed by maltose binding protein (MBP) tag and TEV cleavage site in pET21 vector (Novagen).

SmD1/D2 (or SmD1s/D2) was purified by Ni-column first, followed by TEV protease cleavage, secondary pass of Ni-column, cation exchange, and gel filtration chromatography. SmF/E was purified by a similar procedure except that anion exchange was used instead. SmF/E and SmG were coexpressed and purified in the same way as SmF/E. Gemin2 and SMN_Ge2BD_ were coexpressed and purified by Ni-column first, followed by TEV protease cleavage, Ni-column, and anion exchange chromatography.

To make the heptamer of the Gemin2 (or Gemin2ΔN39)-SMN_Ge2BD_-5Sm complex, equal molar amount of the SmD1s/D2, SmF/E/G, and Gemin2(or Gemin2ΔN39)/SMN_Ge2BD_ complexes were mixed in gel filtration buffer (20 mM Tris-HCl [pH 8.0], 150 mM NaCl, 1 mM EDTA, and 1 mM TCEP [tris(2-carboxyethyl) phosphine]) supplemented with 0.5 M NaCl, and subjected to superdex200 GFC (HiLoad 16/600 or Increase 10/300 GL, GE Healthcare Bio-Sciences, Sweden). The fractions containing all seven components were checked by SDS-PAGE, pooled and concentrated to 7–12 mg/ml, and used for crystallization studies. To make the hexamer of the Gemin2ΔN39-SMN_Ge2BD_-D1s/D2/F/E complex, equal molar amount of the SmD1s/D2, SmF/E, and Gemin2ΔN39/SMN_Ge2BD_ complexes were mixed in the same gel filtration buffer as above, and subjected to superdex200 GFC. The fractions containing all six components were checked by SDS-PAGE, pooled and concentrated to 4-5 mg/ml, and used for crystallization studies.

### Crystallization, Data Collection and Structure Determination

Human Gemin2-SMN_Ge2BD_-D1s/D2/F/E/G complex (Complex A) crystals were grown in 6% PEG8000, 100 mM Tris-HCl (pH 7.5–8.2), human Gemin2ΔN39-SMN_Ge2BD_-D1s/D2/F/E complex (Complex B) crystals were grown in 4% PEG8000, 100 mM Tris-HCl (pH 7.5–8.2), and human Gemin2ΔN39-SMN_Ge2BD_-D1s/D2/F/E/G complex (Complex C) crystals were grown in 1% PEG8000, 100 mM Tris-HCl (pH 7.5–8.2). They were all grown by hanging-drop vapor diffusion method at 20°C within a couple of days. They all form in space group P2_1_2_1_2_1_, but with various unit cell parameters (Table S1). The crystals were cryoprotected by gradual transfer from reservoir solution containing 10% to 40% PEG400, and frozen in liquid nitrogen. The X-ray diffraction data sets of these complex crystals were collected at beamlines BL17U1 and BL19U1 at the National Facility for Protein Science (NFPS) and Shanghai Synchrotron Radiation Facility (Shanghai, China) at wavelengths of 0.97853 and 0.97846 Å. Data were processed by HKL2000 (46). Since the diffraction of the crystals was severely anisotropic, the data sets were reprocessed and truncated ellipsoidally by anisoscaling(47). The structures were solved by molecular replacement with the 2.5 Å crystal structure (PDB code 3S6N)(14) as the search model by PHASER (48) from CCP4 suite (49). The models were improved by cycles of manual rebuilding in Coot (50) and REFMAC refinement (51). The final data collection and refinement statistics are summarized in Table S1. The coordinates and structural factors of the three complexes, A-C, have been deposited in the Protein Data Bank under ID codes 5XJQ, 5XJS and 5XJR, respectively.

### *In vitro* RNA production and purification

With the exception of the nonameric Sm site, AAUUUUUGA, which was chemical synthesized by Takara, all RNAs, including U4, flU4 and their derivatives (Their sequences and predicted secondary structures are in Table S2 and Fig. S4a) were produced by in vitro transcription using MEGAscript kit (Ambion). The templates were made by either annealing of two complementary primers or PCR. Transcribed RNAs were separated by urea-PAGE and the gel containing the RNAs was cut and collected. RNAs were purified by phenol-chloroform extraction, followed by precipitation using ethanol. After spin-vacuum drying, the purified RNAs were dissolved in buffer containing 20mM Tris-HCl, 250 mM KCl, 2mM MgCl_2_, pH7.5. The qualities of these RNAs were checked on agarose gel electrophoresis (Fig. S4b).

### *In vitro* RNA binding, electrophoresis mobility shift assay

Binding of 7S or 7SΔN complex to U4 or U4ΔSm RNA was performed in total volume of 20 μl solution containing 20mM Tris-HCl (pH 8.0), 250mM NaCl, 2mM MgCl_2_, 1 mM EDTA, and 1mM DTT. Various amounts of reconstituted 7S or 7SΔN complex (2.5, 5, and 10 pM of each) were incubated with 50 pM of U4 or U4ΔSm RNA at 37°C for 40 min. After that, 1/10 (v/v) glycerol was added to the reaction mixture and the RNPs were analyzed by 0.4% native agarose gel electrophoresis. RNA was visualized by SYBR green (Thermo Fisher Scientific).

### *In vitro* RNA-Protein complex assembly assay

RNA-protein complex assembly assays were performed by incubating 5Sm or 7SΔN with various RNAs in final volume of 500μl in assembly buffer containing 20mM Tris-HCl (pH 7.5), 250mM NaCl, 2mM MgCl_2_, 1 mM EDTA, and 1mM DTT, with their amounts described in detail in Table S3 (control proteins or RNAs followed the same procedure). RNAs were pre-incubated at 65°C for 10 min followed by cool-down in room temperature before mixing with proteins. After incubation at 37°C for 40 min, the samples were collected at 15,000 rpm for 5 min in a table centrifuge and applied into superdex200 Increase 10/300 GL GFC via a 500 μl sample loop. The eluted fractions were collected each 0.5 ml, resolved by SDS-PAGE directly (visualized by silver staining) or after concentration to 50 μl (visualized by CBB staining). In certain situations, SDS-PAGE was also stained by nucleic acids dye, Super GelRed (US Everbrigh) and visualized under UV to identify the positions of RNAs. Their GFC positions are summarized in Table S4.

### Simulation of GFC traces of component peaks and the combined traces

The traces of component peaks in GFC can be considered as a Gaussian distribution: 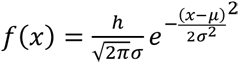. μ is the position (or elution volume) of the peak, σ is the diffusion parameter, positively related to the elution volume, and *h* is the OD value of the peak. For the traces of protein, RNA and RNA-protein complex, OD_260nm_ values are about a half, 2 times and 1.9 times of OD_280nm_ values respectively. For the GFC traces, which contain two or more component peaks, make μ equal to the position of each component peak and adjust *h* and σ (a peak eluted at larger volume has an equal or higher σ value) to fit both the OD_260nm_ and OD_280nm_ traces to be similar to the experimental results by Microsoft Excel software. The separate component peaks are shown together in a plot and the combined trace in a separate plot (Fig.3, Fig. S7, S8). The peak concentration of each component was calculated using its molar extinction coefficient (MEC) (Tables S6-7 and Fig.3, Fig. S7, S8), and the MEC was either calculated by Vector NTI software (Thermo Fisher Scientific) for protein or by the equation: MEC (at 260nm) = 1000 × RNA length/0.11 (Table S5).

## RESULTS

### The narrow conformation of 5Sm bound by Gemin2 is not from crystal packing

In the second phase of Sm core assembly, the crystal structure of the 7S complex we determined previously represents an initial state (14). It reveals how Gemin2 binds 5Sm and SMN. Interestingly, we also observed that the conformation of 5Sm in the 7S complex is narrow compared with the mature snRNP core structures (14,36,38–40)(Fig. 1a-b). As has been mentioned in our previous study (14), although individual Sm proteins between the initial and mature states show little deviations, the width of the opening between SmD1 and SmG in the 7S complex (indicated by the Cα-Cα distance between Asn37 of SmD1 and Asn39 of SmG, the most conserved residues in Sm proteins, being 27.4 Å) is considerably smaller than that in the mature Sm core (the distance is 31.2 Å), and correspondingly, the curvature of the horseshoe-shaped 5Sm in the 7S complex is higher than that in the mature Sm core (Fig. 1a-b). Significantly, the space between SmD1 and SmG in the 7S complex is too narrow for SmD3/B to fit in and the 5Sm’s RNA-binding pocket could not accommodate the snRNA’s Sm site conformation from U1- and U4-snRNP core structures (14). However, it is unknown whether the narrowness of 5Sm in the 7S complex is a real, physiologically relevant state or just an artefact arising from crystal packing, because in the crystal lattice of the 7S complex a second Gemin2’s C-terminal domain (CTD) is located right in between SmD1 and SmG contacting both (Fig. 1c) and crystal packing inducing artificial conformations is well documented (52). It is very likely that the second Gemin2’s CTD pulls SmD1 and SmG close to each other and artificially induces the narrowness of 5Sm. In the first phase of Sm core assembly, there are available structures of two complexes (15), the 6S complex, in which Drosophila pICln binds human 5Sm in a ring shape, and the 8S complex, in which Drosophila Gemin2/SMN-N-terminal domain bind to the peripheral side of 5Sm in the 6S complex. In both these complexes, the conformations of 5Sm are also narrow, and more precisely, even narrower than that in the 7S complex, as indicated by the Cα-Cα distances between Asn37 of SmD1 and Asn39 of SmG being 25.5 Å in the 6S complex and 26.2 Å in the 8S complex, versus 27.4 Å in the 7S complex. The narrowness of 5Sm in both these complexes is because the narrow-sized pICln (occupying only the angular space of one and a half Sm proteins) contacts SmD1 and SmG (15), and therefore cannot provide any clue for the conformation of 5Sm in the 7S complex where pICln is absent. Moreover, the interfaces between Sm and Sm-like proteins are relatively flexible because they can form hexamers, heptamers and even octamers (53–55). It is not implausible that 5Sm bound by Gemin2 in the 7S complex before snRNAs’ assembly is in a wide conformation as it is in the final Sm core (Fig. 1d). So, to study the mechanism of Sm core assembly in the second phase, it is necessary to identify at first whether the conformation of 5Sm bound by Gemin2 is narrow or wide.

**Figure 1.**
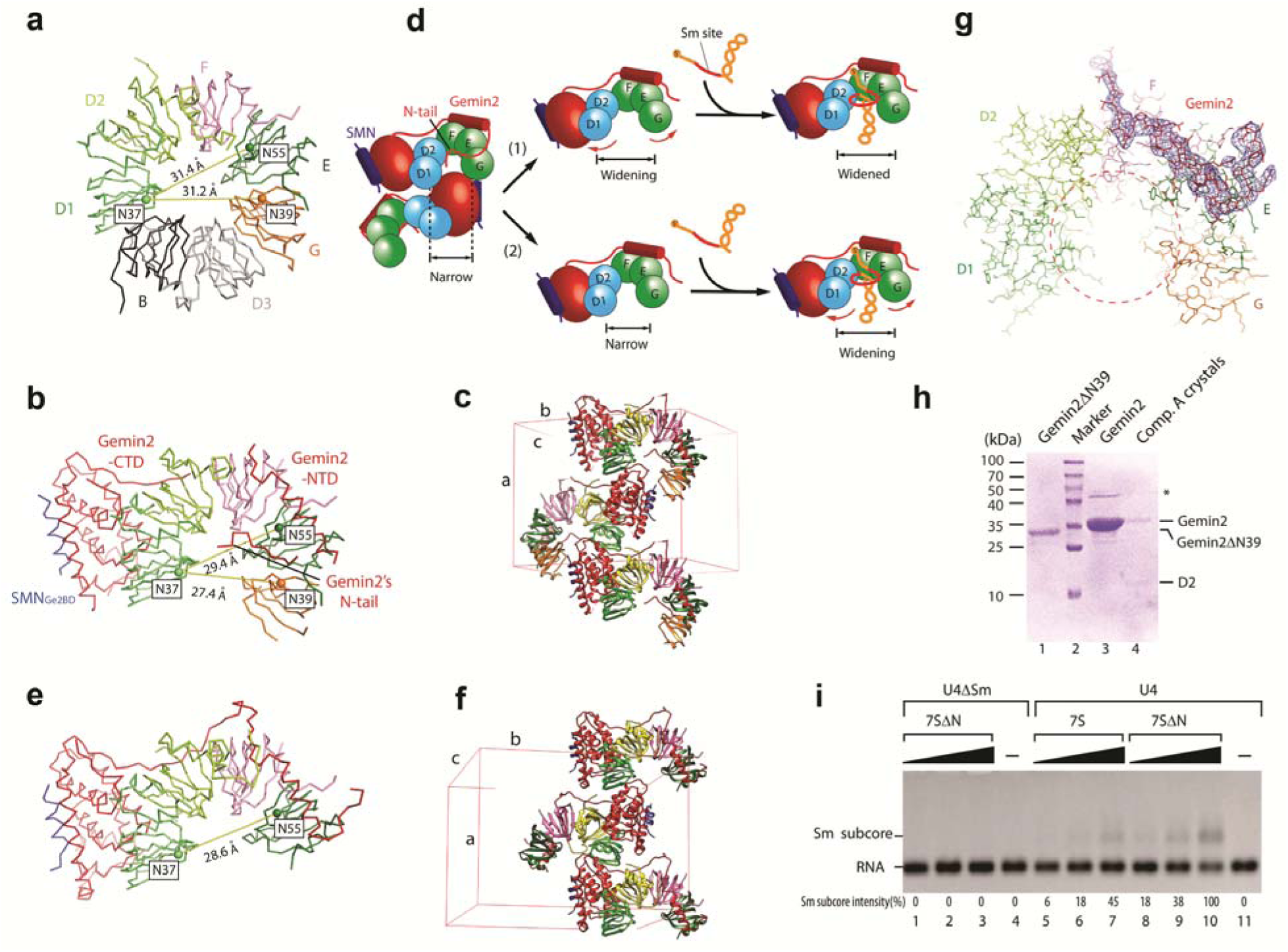
The narrow conformation of Sm heteropentamer bound by Gemin2/SMN_Ge2BD_ is not caused by crystal packing as well as Gemin2’s N-tail. (a) In the mature Sm core represented by U1 snRNP (PDB code 4PJO)(39), the heteropentamer SmD1/D2/F/E/G is in a wide conformation, as indicated by the Cα-Cα distances between N37.SmD1 and N39.SmG, and between N37.SmD1 and N55.SmE. (b) In the previous 7S complex (PDB code 3S6N)(14), the Sm heteropentamer is in a narrow conformation. (c) In the crystal lattice, the opening between SmD1 and SmG of one 5Sm is occupied by Gemin2’s CTD of a second complex, which may induce the narrowness of 5Sm. (d) there are two possible mechanisms of the assembly of snRNA into the Sm subcore bound by Gemin2/SMN_Ge2BD_. Without the influence of crystal packing and Gemin2’s N-tail inside 5Sm, 5Sm either automatically becomes widened before snRNA’s binding (1), or still keeps the narrow conformation until snRNA’s assembly (2). (e) In the structure of Complex B, which has no SmG and no Gemin2’s N-tail, the conformation of the Sm proteins is still narrow. (f) In the crystal lattice of Complex B, only SmD1 contacts Gemin2’s CTD of a second complex. (g) Gemin2’s N-tail is outside the RNA-binding pocket in the crystal structure of Complex A, in contrast to the previous 7S structure (PDB code 3S6N)(14). Only Gemin2’s N-terminus and 5Sm are shown. SigmaA-weighted 2Fo-Fc electron density maps (blue meshes) are contoured at 1.1σ of Gemin2 N-terminus (residues 1-76). Red circles indicate the RNA-binding pocket. The seven Sm proteins, D1, D2, F, E, G, D3 and B, are colored in green, lemon, pink, dark green, orange, light grey and dark grey respectively. Gemin2 and SMN_Ge2BD_ are colored in red and blue respectively. Unit cells for the previous 7S complex and Complex C are showed in (c) and (f). (h) The Gemin2 in Complex A crystals was intact. Crystals of complex A were picked up and subjected to SDS-PAGE and CBB staining. Full-length Gemin2 and Gemin2ΔN39 were used as controls. The asterisk indicates impurity. (i) Gemin2’s N-tail accounts for only about 2-fold of U4 RNA inhibition to Gemin2ΔN39. Three concentrations (2.5, 5 and 10 pM) of reconstituted 7S or 7SΔN complex were pre-incubated with U4 or U4ΔSm snRNA at 37°C for 40 min and subjected to electrophoresis mobility shift assay. One representative from three independent results is shown. The positions of the free RNA and 5Sm assembled on the RNA are indicated. The levels of Sm subcore were quantitated using imaging software and normalized. See also Fig. S1-2.

To test this, we used a crystallographic approach and attempted to pack the 7S complex and its derivatives in different crystal lattices to avoid the above packing contacts. After we failed at obtaining a different crystal of the original 7S complex by trying different solution conditions, we reconstituted Complex A, a 7S complex with a short version of SmD1, SmD1s, in which the C-terminus (residues 83-119, which is invisible in the previous 7S complex structure (14)) was truncated, replacing SmD1. Complex A formed crystals in different crystal lattice and its packing is significantly different from the previous 7S complex (Table S1 and Fig. S1), however, Gemin2’s CTD is still located in between the SmD1 and SmG of another complex and the distance between the SmD1 and SmG is little altered. We also reconstituted several other 7S complex variations, among which the most significant one is Complex B, a derivative of Complex A without SmG and with Gemin2ΔN39 (the N-terminal residues 1-39 are truncated to further test the effect of the N-tail) replacing Gemin2 (Fig. 1e-f, S1 and Table S1). In the crystal lattice, although SmD1 is still in contact with Gemin2’s CTD of a second complex, due to the absence of SmG, SmE at the other end of the crescent Sm hetero-oligomer is far away from the second Gemin2’s CTD for interaction, eliminating the influence of crystal packing on the curvature of the Sm hetero-oligomer (Fig. 1f). However, the curvature of D1/D2/F/E is little different from that of the original complex as indicated by no increase of the Cα-Cα distance between the most conserved residues Asn37 of SmD1 and Asn55 of SmE (28.6 Å in Complex B vs. 29.4 Å in the previous 7S complex) (Fig. 1b,e). So, this observation indicates that the narrow conformation of 5Sm bound by Gemin2 is not caused by crystal packing artifact, but is a physiologically relevant one. We term this conformation as the ground state.

### The narrowness of 5Sm is not caused by Gemin2’s N-terminal tail

Since in the previous structure of the 7S complex the N-terminal tail (N-tail, residues 22-31 visible) of Gemin2 is located inside the central RNA-binding pocket, it is possible that this N-tail induces the narrowness of the 5Sm. To test this possibility, we reconstituted Complex C, a derivative of Complex A with Gemin2ΔN39 replacing Gemin2, for crystallization. The crystal of Complex C had the same space group as Complexes A and B and similar unit cell parameters to them (Table S1). However, the curvature of 5Sm is still the same as that of the previous complex (Data not shown). In addition, we also created Complex B (see above) for crystallization. The absence of Gemin2’s N-tail did not change the curvature of D1/D2/F/E (Fig. 1e-f). These data demonstrate that Gemin2’s N-tail does not play a role in the narrowness of 5Sm and the narrowness of 5Sm is caused by the rest part (residues 40-280) of Gemin2.

### Gemin2’s N-tail flips dynamically and plays a minor inhibitory role in RNA binding

In addition, surprisingly, in one 7S complex crystal structure with the full-length Gemin2 (Complex A), we observed that there was no electron density inside the RNA-binding pocket of 5Sm (Fig. 1g, Table S1 and Fig. S1), in contrast to the previous complex structure where Gemin2’s N-tail is inside the RNA-binding pocket (14). Complex A was crystallized under a condition similar to the previous 7S complex, but has a different crystal packing (Table S1 and Fig. S1). Checking the components of crystal sample by SDS-PAGE, we saw the band of Gemin2 keeping the original full-length size, indicating no degradation (Fig. 1h). So it is only reasonable to explain that Gemin2’s N-tail was located outside and flexible in the crystal of Complex A. These data indicate that Gemin2’s N-tail may not be located firmly in one place; instead its positions may be quite dynamic. Consistent with this was that the peak of 7S (containing the full-length Gemin2) eluted earlier than SMN(26-62)/Gemin2ΔN39/5Sm (7SΔN) in gel filtration chromatography (GFC) (Fig. S2), indicating that Gemin2’s N-tail flips outside the pocket and increase the complex’s size.

In the previous study, an inhibitory role of Gemin2’s N-tail on snRNAs’ binding to 5Sm was observed (14). However, the experiments were carried out by mixing separate SMN_Ge2BD_/Gemin2 (or SMN_Ge2BD_/Gemin2ΔN39) and 5Sm at various ratios, and this way could not faithfully mimic the physiological state, in which 5Sm binds to Gemin2/SMN in an equal stoichiometry. In this study, we used preformed 7S and 7SΔN to examine snRNA binding with the representative U4 snRNA, which has features matching the snRNP code and had proved to assemble into the Sm core (56). Using electrophoresis mobility shift assay (EMSA), we observed that 7S or 7SΔN formed Sm subcore with U4 snRNA as its concentration increases, while the highest concentration of 7SΔN tested could not bind the negative control, U4ΔSm RNA, in which the nonameric Sm site was replaced by AACCCCCGA (Fig. 1i). 7SΔN formed more subcore with U4 snRNA than 7S, but its binding efficiency was only about 2-fold higher than that of 7S (Fig. 1i). As U4 snRNA has the same snRNA code as the other snRNAs (9), we concluded that Gemin2’s N-tail has a minor inhibitory role in 7S binding to snRNAs. This conclusion is consistent with the crystallographic and chromatographic observations and the dynamic nature of Gemin2’s N-tail.

### 5Sm in 7SΔN cannot bind RNA containing the Sm site only or at 3’-end

In *in vitro* experiments, mixing D1/D2, F/E/G and a 9-nucleotide Sm-site RNA (9nt), AAUUUUUGA, could readily produce a stable Sm subcore (6). We wondered whether a preformed 7S with 5Sm in the narrow state would similarly accept a Sm-site RNA. Because Gemin2’s N-tail contributes little to Gemin2’s binding to 5Sm and to the conformation of 5Sm, but its minor inhibitory role in snRNA binding could interfere with evaluation of the narrow conformation of 5Sm in RNA binding, to simplify our analysis, we used Gemin2ΔN39 instead of the full-length Gemin2 to perform RNA binding experiments. Furthermore, to facilitate analysis of complex components by taking advantage of purified proteins and RNAs, we adopted GFC plus SDS-PAGE analysis (two-dimensional separation) instead of EMSA (one-dimensional separation). Three parameters were generally monitored for the formation of a RNA-protein complex: peak elution volume (position), ratio of OD_260nm_ to OD_280nm_ (OD_260/280_), which can distinguish protein (OD_260/280_ is about ½), RNA (OD_260/280_ is about 2) and protein-RNA complex (OD_260/280_ is between 1 and 2 and close to 2 because RNA dominates OD values), and component analysis by SDS-PAGE followed by silver staining or Coomassie brilliant blue (CBB) staining. Whenever the GFC traces were complicated, simulations of the separate and the combined peaks were provided (Fig.3 & S7-8). Moreover, the molar concentrations of the separate peaks were calculated (Tables S4-7). Using the 9nt, AAUUUUUGA, to incubate with D1/D2 and F/E/G followed by GFC separation, we observed the formation of Sm subcore (the peak at 14.37 ml with OD_260/280_ of 1.12) (Fig. S3, a-d, compare panel d with a-c), which is consistent with the early report (6). To better detect RNA by SDS-PAGE and silver staining, we used a longer RNA (37nt) containing the Sm site at its 3’ end (3’ Sm) to perform the same experiment and observed the formation of Sm subcore (the peak at 13.51 ml with OD_260/280_ of 1.89) (Fig. 2, a-c, compare panel c with a-b). Surprisingly, however, using the same 3’ Sm RNA to incubate with the preformed 7SΔN complex, we could only see the 7SΔN peak (13.78 ml) but no formation of any RNA-protein complex (Fig. 2, d-f, compare f with a, d and e). This indicated that the Sm-site RNA, even with additional single-stranded RNA at its 5’-end, cannot bind to 5Sm when the latter is bound by Gemin2ΔN39/SMN_Ge2BD_. These results suggest that the narrow conformation of 5Sm bound by Gemin2 plays a restrictive role in RNA binding.

**Figure 2.**
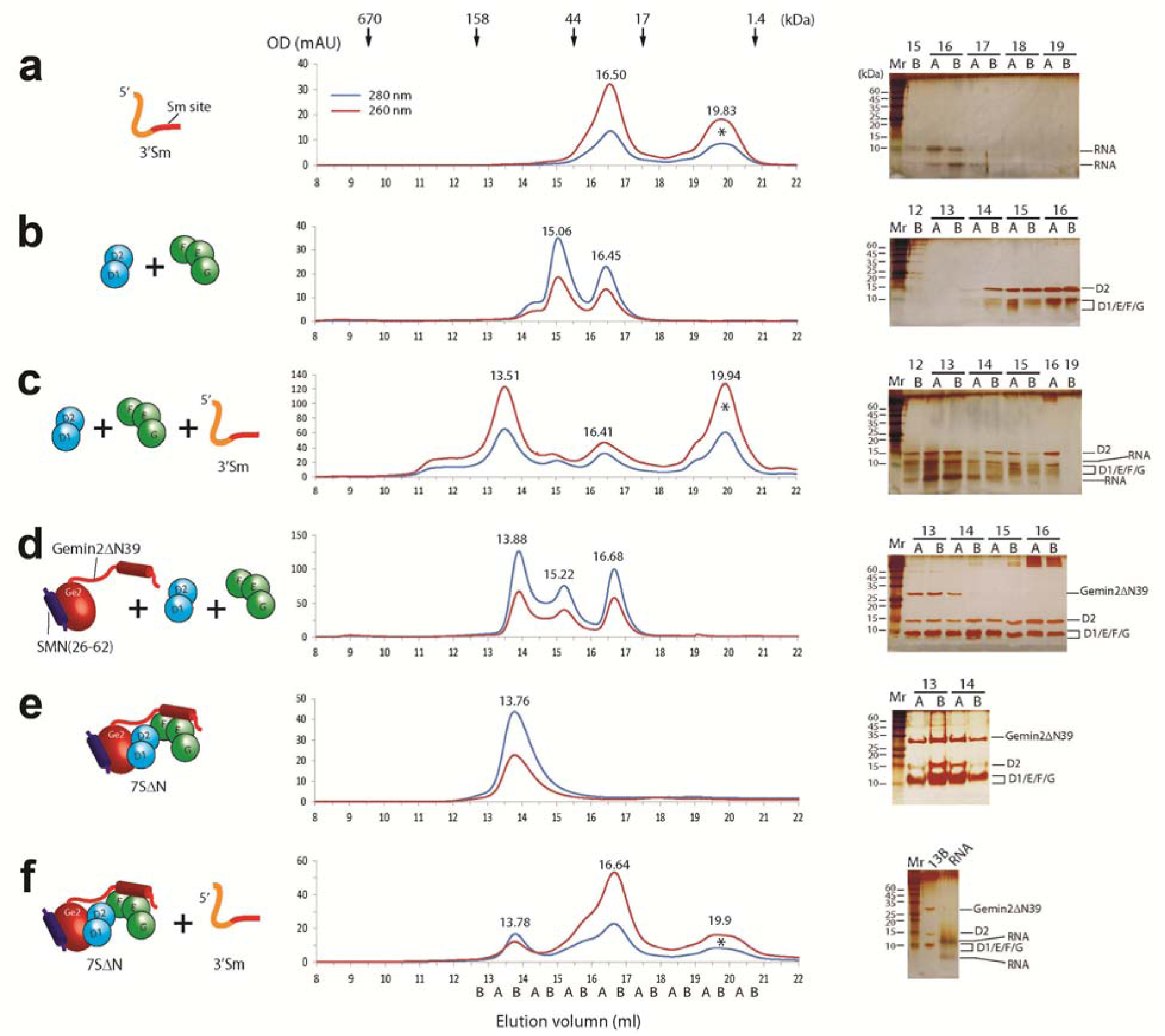
The 7SΔN complex cannot bind RNA containing the Sm site at 3’ side. Different components (in cartoon, left panels) were subjected to GFC analysis (middle panels) followed by SDS-PAGE plus silver staining (right panels). Individual fractions were collected half a milliliter per fraction and named on the basis of elution position with A and B as the first and second half ml. The input components were (a)3’Sm, (b)D1/D2 and F/E/G, (c) 3’Sm, D1/D2 and F/E/G, (d)SMN(26-62)/Gemin2ΔN39 and extra amounts of both D1/D2 and F/E/G, (e)The fraction of the 7SΔN peak at 13.88 ml from (d), and (f) 7SΔN and 3’Sm. For each, one representative result from at least two independent experiments is shown. The small-sized SMN(26-62) peptide was invisible on SDS-PAGE/staining but must be present as can be seen in the crystal structures of the 7S complex and mutants. See also Fig. S3.

**Figure 3.**
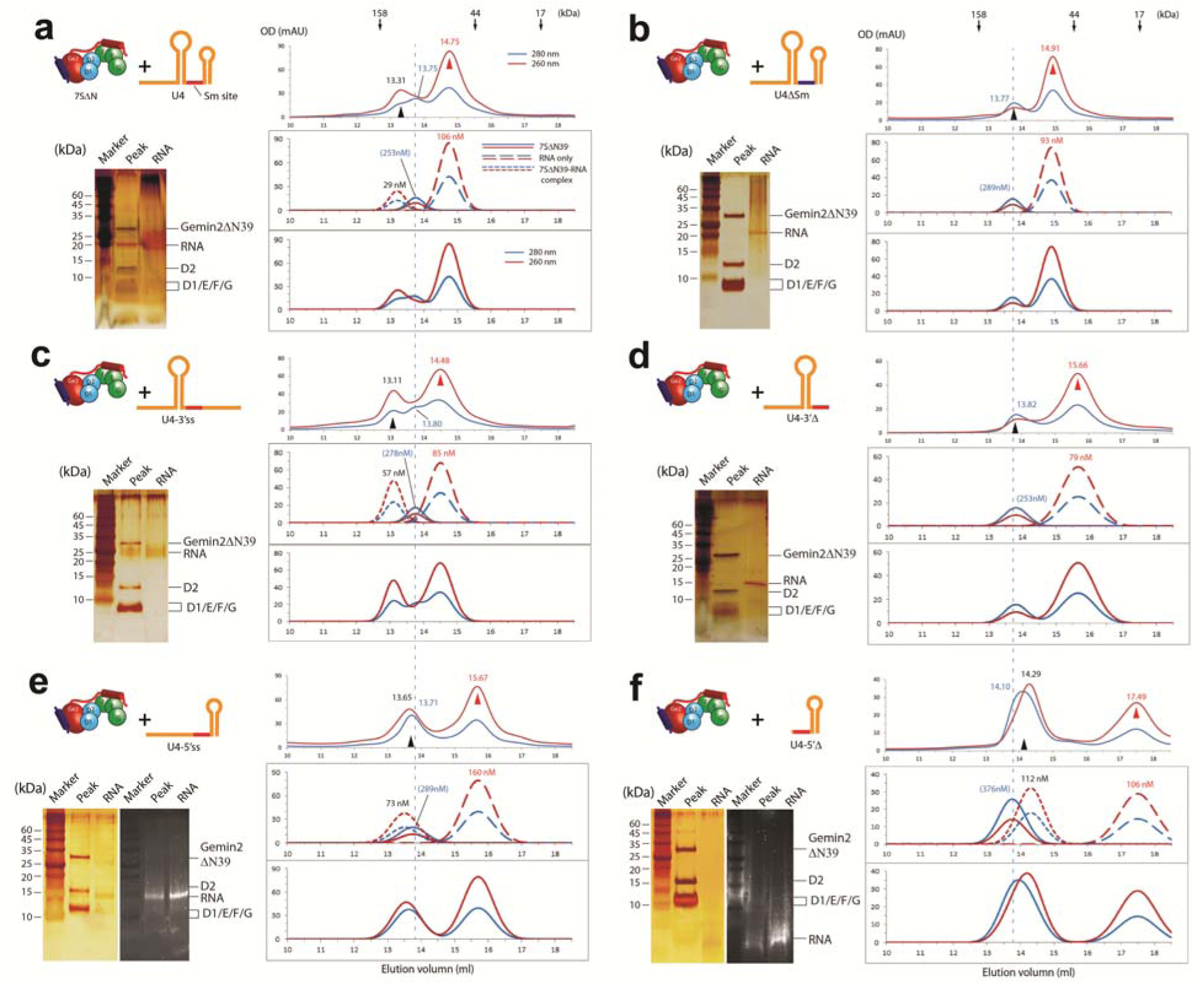
The 7SΔN complex selectively binds RNAs with both the Sm site and a 3’ RNA. 7SΔN was pre-incubated with various RNAs (in cartoon, left top in each panel): U4 (a), U4ΔSm (b), U4-3’ss (c), U4-3’Δ (d), U4-5’ss (e) or U4-5’Δ (f), and the mixture was subjected to GFC (right top in each panel). The front peak fractions (pointed at by black arrow heads) and RNA inputs were analyzed by SDS-PAGE plus silver staining (left bottom in each panel). For (e) and (f), additional staining by nucleic acids dye, Super GelRed (US Everbrigh), was used on duplicated gels to identify the positions of RNAs. For each, one representative result from three independent experiments is shown. The peaks pointed at by the red arrow heads are the RNA peaks. The blue vertical dash lines indicate the peak position of 7SΔN. The simulations of the component peaks and their concentrations (see Table S6 for the calculation. The values without brackets are more accurate than those in brackets) and the combined traces are shown (right bottom in each panel). See also Fig. S4-5.

### RNA binding to 5Sm in 7SΔN needs both the Sm site and its 3’-sequence

This surprising observation triggered us to ask what RNA feature 7SΔN can recognize. Does it match the snRNP code previously identified by cell-based experiments (9)? All the RNAs, including snRNAs and a few viral RNAs, which can be assembled into snRNP in cells, contain the nonameric Sm site with two SLs flanking it (9,29,56). Previous studies have showed that the nonameric Sm site alone is both essential and sufficient for Sm core assembly *in vitro* (6), but for *in vivo* assembly, an additional feature, the 3’SL, is also required. The Sm site and the 3’SL together are termed as the snRNP code (9). In the crystal structures of the fully-assembled U1-snRNP and U4-snRNP cores (39, 40), the Sm site and the two flanking SLs all have contact with the Sm proteins. The first seven bases of the nanomeric Sm site form a circular plane and specifically interact with the seven Sm proteins inside the upper portion of the central channel, the phosphate groups of the 3’-SL interact nonspecifically with the lower portion of the central channel, and the 5’SL contacts the outside of the Sm hepatomeric ring.

So, we decided to systematically study the Sm site and both SLs of snRNAs in binding to 7SΔN. We used a representative snRNA, a middle-sized, 3’ fraction of human U4 snRNA (U4 snRNA), which has SLs flanking the Sm site (containing all the features described above) and had proved to assemble into the Sm core (56), and designed its several derivatives (their sequences, secondary structures and qualities are in Table S2 and Fig. S4) by mutating the Sm site, linearizing or deleting the SL at either side of the Sm site, one at a time, to test their binding to the preformed 7SΔN via the same procedure as above. As expected, using the U4 snRNA to incubate with 7SΔN, we observed the formation of Sm subcore (peak at 13.31 ml with OD_260/280_ of 1.9) from GFC separation (Fig. 3a). However, Sm subcore only formed in a small percentage, because there were still free 7SΔN and free RNA (their peaks at 13.75ml and 14.75ml respectively) in substantial amount (Fig. 3a, compare the experimental trace with the simulation). In contract, using U4ΔSm snRNA, we only observed the peak of 7SΔN (at 13.77 ml with OD_260/280_ of 0.74) at the front, indicating no formation of Sm subcore on U4ΔSm snRNA (Fig. 3b). These results are consistent with the EMSA result (Fig.1i). These observations indicate that the narrow conformation of 5Sm bound by Gemin2 plays a restrictive role in RNA binding, for the 7SΔN state not only cannot bind RNA containing the Sm-site only or at 3’-side, but can only bind the normal snRNA containing the snRNP code to a limited degree.

Next, we tested the 3’-side of U4 RNA on binding to 7SΔN. We changed the 3’-SL of U4 snRNA to a linear single strand (U4-3’ss). The GFC trace showed three peaks eluted at 13.11, 13.80 and 14.48 ml (Fig. 3c), which were the 7SΔN-RNA complex, 7SΔN and RNA respectively. This observation suggests that single-stranded RNA at the 3’ end of the Sm site can still bind to 7SΔN to a small degree. When the 3’SL was completely removed (U4-3’Δ), however, the RNA did not bind to 7SΔN as showed by the absence of an earlier peak than the 7SΔN peak (at 13.82 ml) (Fig. 3d). This result indicates that the presence of RNA at the 3’ side of the Sm site, either single- or double-stranded, is critical for the formation of a 7SΔN-RNA complex. If there is no RNA at the 3’ side, any RNA, either a single- or double-stranded, at the 5’ side of the Sm site cannot make the RNA bind to 7SΔN.

Finally, we tested the 5’-side of U4 RNA on binding to 7SΔN. When we linearized the 5’-SL of the U4 snRNA (U4-5’ss), or deleted the 5’SL of U4 snRNA (U4-5’Δ), we observed an additional peak of RNA-containing complex (13.65 ml or 14.29 ml, because of its OD_260/280_ >1 and the presence of RNA in the peak by silver staining) eluted much earlier than the RNA peak (15.67 ml or 17.49 ml) (Fig. 3, e-f). These observations indicate that whether the 5’ side of the Sm site is single- or double- stranded or absent, as long as RNA contains the Sm site plus 3’-SL, it can bind to 7SΔN. In summary, these observations showed that the 5’-side RNA of the Sm site is not required for RNA binding, but a single-stranded RNA at the 3’ side of the Sm site seems necessary and sufficient to bind to 7SΔN. To further confirm this conclusion, we used a minimal RNA containing only the Sm site and 3’ single-stranded RNA (U4-5’Δ-3’ss) to perform the assay. As we expected, an RNA-protein complex formed at 13.97 ml, which is much earlier than the RNA alone (16.62 ml) (Fig. S5f).

### Gemin2 starts to be released during Sm subcore assembly

In the above 7SΔN binding test with U4-5’ss or U4-5’Δ RNA, both of which have the absence of a 5’-adjacent SL at the Sm site, however, we also observed a surprising phenomenon at the early RNA-containing peak, that is, the difference of the OD_260_ and OD_280_ peak positions (13.65 and 13.71 ml respectively in Fig. 3e, and 14.29 and 14.10 ml in Fig. 3f). This indicated that there should be two peaks of similar sizes eluted, one being 7SΔN (around 13.8 ml, and with lower ratio of OD_260/280_), and the other being a protein complex containing RNA (Fig. 3e-f. compare the experimental traces with the simulated ones). But the identity of the latter complex was perplexing, especially for that containing U4-5’Δ RNA (Fig. 3f), which strikingly eluted later than 7SΔN. If the complex was 7SΔN plus RNA, the size should be larger than 7SΔN and should be eluted no later than 7SΔN alone. It must not be RNA alone because RNA alone eluted only at 17.48 ml (Fig. 3f. and Fig.S5c). At this point, there would be two possibilities: either the binding of RNA to 7SΔN changes the conformation and reduces its hydrodynamic radius or there was a loss of protein components upon the binding of RNA to 7SΔN, logically Gemin2/SMN_Ge2BD_. Although the latter was buttressed by further experiments in which the Sm subcores reconstituted from these RNAs with D1/D2 and F/E/G were eluted at similar positions to their corresponding OD_260_ peaks in binding to 7SΔN described above (Fig. S5a-e), due to the overlapping of these RNA-containing protein complexes with 7SΔN at GFC, at this point we were unable to make a clear distinction.

To better monitor whether Gemin2ΔN39/SMN_Ge2BD_ is released, to which extent, and at which step of Sm-core assembly, we created large-sized, full-length U4 snRNA (flU4) and its several derivatives (their sequences, secondary structures and qualities are in Table S2 and Fig. S4; their GFC traces alone are in Fig. S6), the assembly of which into Sm subcores and cores would potentially elute earlier and have a better separation from 7SΔN as well as Gemin2ΔN39/SMN_Ge2BD._ In addition, from the above experiments, we noticed that three types of RNAs, (1) like U4, in which 2 SLs tightly flank the Sm site, (2) like U4-5’ss or U4-5’Δ, containing the Sm site plus a 3’SL but no 5’SL, and (3) like U4-5’Δ-3’ss, containing the Sm site plus a 3’ single strand but no 5’SL, were able to bind to 7SΔN, but they seemed to behave differently in terms of their expected Sm subcore sizes and protein components (Fig.3 a, c, e-f). Therefore, we used the large-sized RNAs corresponding to the above three types of RNAs in the following experiments to examine the release of Gemin2ΔN39/SMN_Ge2BD_.

At first, the negative control, flU4ΔSm RNA, incubated with 7SΔN for GFC, had no RNA-containing complex formed, but only the separate RNA peak (12.93 ml) and 7SΔN peak (13.75 ml) (Fig. 4a, compare it with Fig. 2e and the simulation in Fig. S7a). In contrast, incubating flU4 snRNA (type 1 RNA) with 7SΔN, we observed that a small peak containing RNA-protein complex appeared the earliest at 12.26 ml (Fig. 4b, compare it with 4a and S7b). This peak fraction contained all 5 Sm proteins and Gemin2ΔN39, but the stoichiometry of Gemin2ΔN39 to 5Sm was less than 1:1 (Fig. 4b. compare lanes 11B and 14A). These experiments showed that Gemin2ΔN39/SMN_Ge2BD_ have started to dissociate from 5Sm when flU4 snRNA binds to 5Sm.

**Figure 4.**
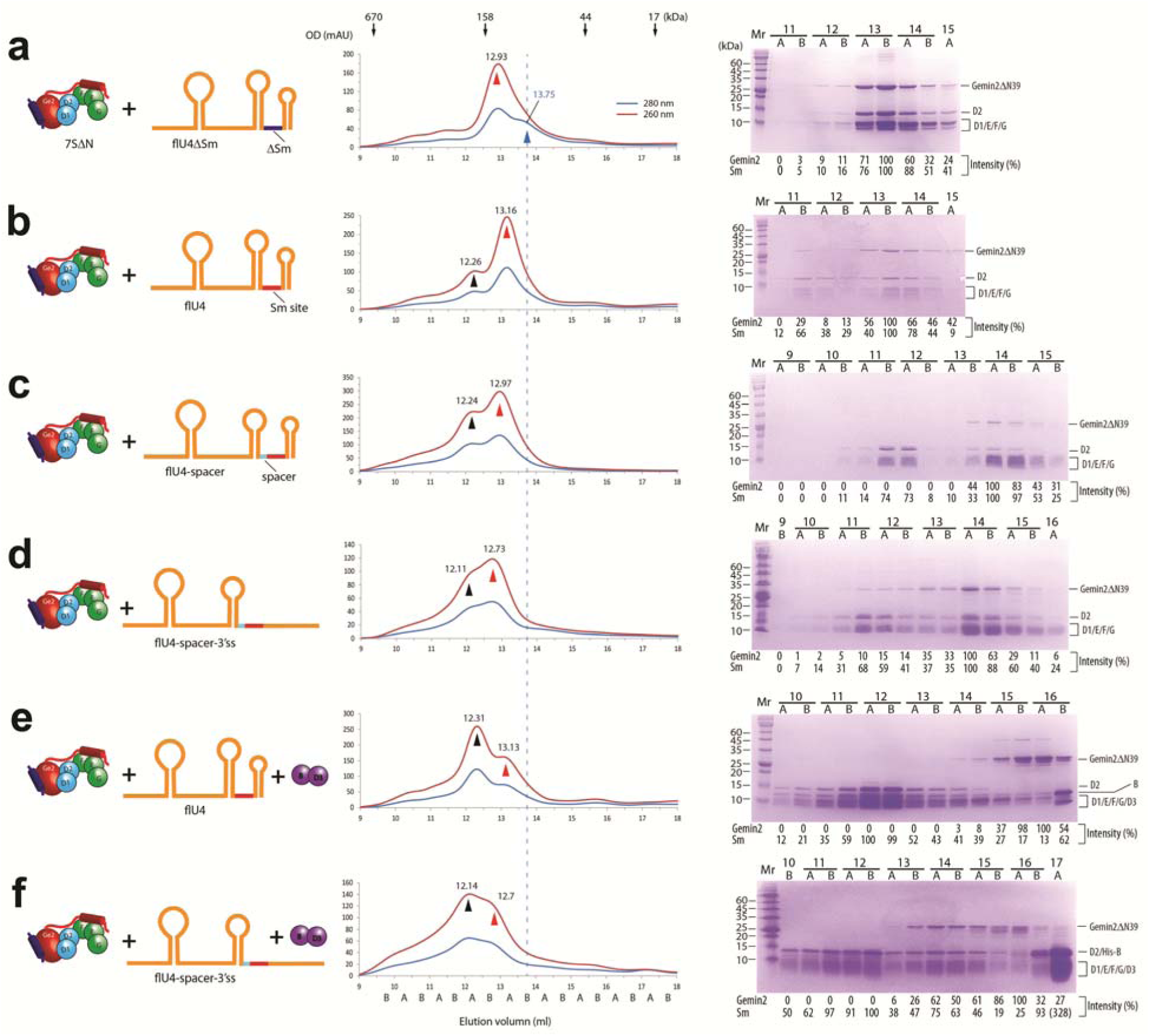
Release of Gemin2 during the assembly of Sm subcores and cores. 7SΔN was pre-incubated with various RNAs (in cartoon, left panels): flU4ΔSm (a), flU4 (b), flU4-spacer (c) or flU4-spacer-3’ss (d) for Sm subcore assembly, or with D3(1-75)/B(1-91) and flU4 (e) or flU4-spacer-3’ss (f) for Sm core assembly, and the mixture was subjected to GFC (middle panels) followed by SDS-PAGE plus CBB staining (right panels). For each, one representative result from at least two independent experiments is shown. CBB staining was chose here for better protein component analysis. Individual fractions are named on elution position, with A and B indicating the first and second half ml. The peaks pointed at by the red arrow heads are the RNA peaks, and the peaks pointed at by the black arrow heads are the Sm-RNA complexes (subcores and cores). The vertical dash line and the blue arrow head indicate the peak position of 7SΔN. The relative intensities of Gemin2ΔN39 and Sm proteins were quantitated using imaging software and normalized. The GFC traces of these RNAs alone are in Fig. S6. The simulations of the component peaks (and concentrations) and the combined traces for each GFC are shown in Fig. S7.

In the previous section, we tested the binding of type 2 RNA (U4-5’ss or U4-5’Δ) to 7SΔN, and observed an aberrant peak of RNA-protein complex. We suspected that the Sm subcore formation on this type of RNA causes a complete detachment of Gemin2ΔN39/SMN_Ge2BD_. To test it, we made a derivative of flU4 snRNA, flU4-spacer (type 2 RNA), in which a room was created between the Sm site and its 5’ SL by disrupting the bottom 3 base pairs of 5’SL (replacing the 3 nucleotides, GGC, at the 3’-end of the 5’SL, with CCG). The incubation of flU4-spacer with 7SΔN gave rise to a Sm subcore with a complete removal of Gemin2ΔN39 (Fig. 4c, lanes 11A-12A), indicating that a free or SL-free 5’ end of the Sm site did cause a complete release of Gemin2ΔN39/SMN_Ge2BD_ from the Sm subcore. But the presence of free RNA (peak at 12.97 ml, also see Fig. S7c) and free 7SΔN (Fig. 4c, lanes 13B-14B) indicated that the formation of the subcore was in an equilibrium with the reactants.

To test the assembly of the type 3 RNA, we used a further derivative of flU4 snRNA, flU4-spacer-3’ss, which linearized the 3’ SL on the basis of flU4-spacer. The incubation of flU4-spacer-3’ss with 7SΔN generated a gel filtration profile similar to flU4, in which a small fraction of Sm subcore formed (peak at ∼12.11 ml, compare with Fig. S7d), to part of which Gemin2ΔN39 was still bound (Fig. 4d, lanes 11B-12A).

### SmD3/B binds and Gemin2 is released during Sm core assembly

As we have shown, 7SΔN has a narrow conformation of 5Sm, which is in conflict with SmD3/B binding. It is consistent with our previous study, in which SMN/Gemin2 can bind only 5Sm even when all 7 Sm proteins are present (14). Does the binding of RNA to 7SΔN expand the SmD1-G opening to allow SmD3/B to join? What about Gemin2’s release upon Sm-core assembly? To test these, we incubated flU4 snRNA (type 1 RNA), 7SΔN and D3(1-75)/B(1-91) (the nonessential C-terminal tails of both are truncated), and subjected the mixture to GFC. The earliest and also highest peak (12.31 ml) contained RNA and all 7 Sm proteins but no Gemin2ΔN39 (Fig. 4e, lanes 12A-12B). The band of Gemin2ΔN39 appears on the GFC at 15.5-16.5 ml as analyzed by SDS-PAGE (lanes 15B and 16A), consistent with the position of Gemin2ΔN39/SMN_Ge2BD_ alone (peak at ∼15.66 ml, Fig. S3e), indicating that the Gemin2ΔN39/SMN_Ge2BD_ was released and in a free state. Furthermore, few 7SΔN complex components seen around the 7SΔN peak of 13.8 ml (Fig. 4e, lanes 13B and 14A) indicated that almost all SmD1/D2/F/E/G pentamer was assembled into the Sm core. These results showed that Sm-core assembly goes to completion upon the joining of SmD3/B to the 7SΔN-RNA complex, and simultaneously causes a complete release of Gemin2ΔN39/SMN_Ge2BD_. For flU4-spacer (type 2 RNA), which seems more efficient in forming the Sm subcore than flU4 (Fig. 4b-c), the addition of D3/B would also drive the Sm core formation to a completion as in the case of flU4 snRNA.

The incubation of flU4-spacer-3’ss (type 3 RNA) with both 7SΔN and D3(1-75)/B(1-91) also produced the Sm core and caused Gemin2ΔN39 to dissociate, but the assembly did not proceed to a completion, as indicated by the presence of free RNA (peak at ∼12.7 ml), 7SΔN (lanes 13B-14B) and D3/B (lanes 16B-17A) (Fig. 4f). This indicated that type 3 RNAs (Sm site plus a 3’-single strand), although they can assemble into the Sm core, are less efficient substrates than types 1 and 2 RNAs (both contain Sm site plus a 3’-SL).

### RNAs containing the snRNP code assemble into 5Sm of 7SΔN more efficiently

The Sm site plus either a 3’-SL or a single-stranded RNA can bind the 7S complex and assemble into the Sm core. But the above experiments suggested that they might have different efficiency. To directly compare assembly efficiency, we performed a competition study by incubating 7SΔN with equal molar amount of flU4-spacer and U4-5’Δ-3’ss. The large difference of their RNA sizes makes their subcore formation visible on SDS-PAGE. The major fractions of Sm subcore containing flU4-spacer appeared on lanes 11A-12B (relative intensity of 1.0, with peak at ∼12.3 ml), whereas the major fractions of Sm subcore containing U4-5’Δ-3’ss on lanes 13B-15A (relative intensity of 0.6, with peak at ∼14.0 ml) (Fig. 5a, compared with Fig. S5f, also see the simulated traces in Fig. S8). Although the latter overlapped 7SΔN (with peak at ∼13.8 ml), which discouraged precise quantification, a simple comparison of the staining intensity of the Sm proteins clearly showed that the Sm subcore containing flU4-spacer dominated, indicating that the snRNP code is more efficient than the Sm site plus a 3’ single strand in subcore formation. In addition, we swapped the 5’ portions of the RNAs and performed a competition study by incubating 7SΔN with equal molar amount of flU4-spacer-3’ss and U4-5’Δ. This time, the staining intensity of the major fractions of the Sm subcore containing flU4-spacer-3’ss became weak (lanes 11A-12B, relative intensity of 1.0, with peak at ∼12.1 ml), while that of the Sm subcore containing U4-5’Δ became strong (lanes 13B-15A, relative intensity of 3.5, with peak at ∼14.3 ml) (Fig. 5b. also compare panel a with b). This result confirmed that RNAs containing the snRNP code assembles more efficiently than RNAs containing the Sm site plus a 3’ single strand. We also incubated 7SΔN with equal amount of the two RNAs, U4-5’Δ-3’ss and U4-5’Δ, which were identical in length and had no 5’ extra portion, for GFC analysis (Fig. S5g. compare panel g with e-f). Consistent with our anticipation, the front peak of OD_260_ appeared at 14.25 ml, close to the peak of the Sm subcore containing U4-5’Δ (14.29 ml), while the free RNA appeared at 16.88 ml, close to the peak of free U4-5’Δ-3’ss (16.62 ml) instead of the peak of free U4-5’Δ (17.48 ml), indicating that more U4-5’Δ was assembled into the Sm subcore.

**Figure 5.**
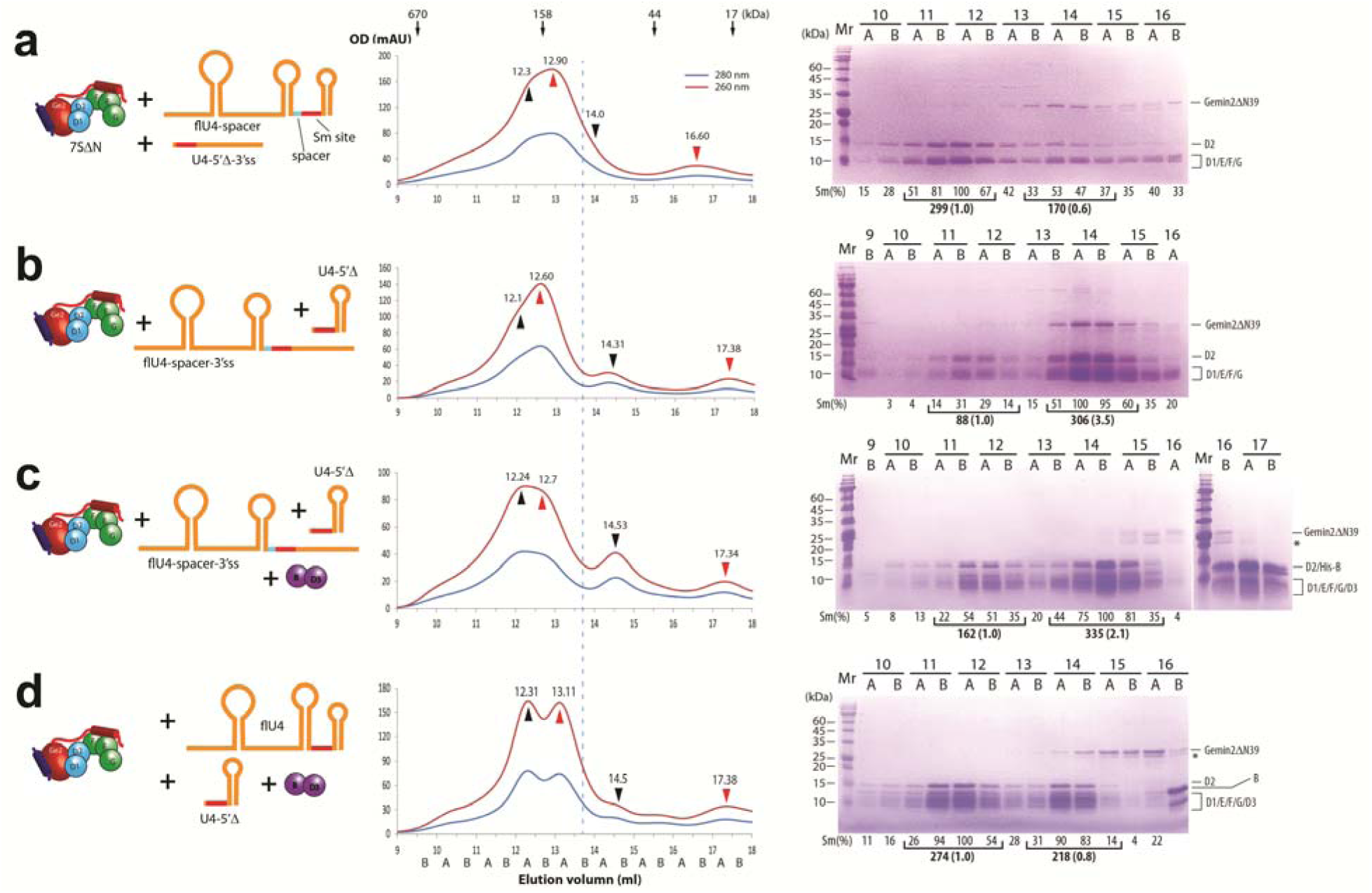
RNAs containing the snRNP code assembles into Sm subcores and cores selectively. 7SΔN was pre-incubated with equal molar amount of flU4-spacer and U4-5’Δ-3’ss (a), or equal molar amount of flU4-spacer-3’ss and U4-5’Δ (b) for Sm subcore assembly. 7SΔN was pre-incubated with D3(1-75)/B(1-91) and equal molar amount of flU4-spacer-3’ss and U4-5’Δ (c) or equal molar amount of flU4 and U4-5’Δ (d) for Sm core assembly. The input components are showed in cartoon (left panels). The mixtures were subjected to GFC (middle panels) followed by SDS-PAGE plus CBB staining (right panels). For each, one representative result from at least two independent experiments is shown. CBB staining was chose for better protein component analysis. Individual fractions are named on elution position, with A and B indicating the first and second half ml. The peaks pointed at by the red arrow heads are the RNA peaks (up, large RNA; down, small RNA), and the peaks pointed at by the black arrow heads are the Sm subcores and cores (up, containing large RNA; down, containing small RNA). The vertical dash line indicates the peak position of 7SΔN. The relative intensities of Sm proteins, Sm(%), were quantitated using imaging software and normalized. The fractions representing the Sm core assembly on large RNA (front) and on small RNAs (back) were combined and further normalized in brackets. The simulations of the component peaks (and their concentrations) and the combined traces for each GFC are shown in Fig. S8.

In addition, to compare the assembly efficiency of the final Sm core, we made a competition analysis by incubating equal molar amount of flU4-spacer-3’ss and U4-5’Δ with both 7SΔN and D3(1-75)/B(1-91). The peak of the Sm core containing flU4-spacer-3’ss appeared at about 12.2 ml, whereas the peak of the Sm core containing U4-5’Δ at about 14.5 ml (Fig. 5c). Gemin2ΔN39 came later, with the darkest band in lane 15B (15.5-16.0 ml, Fig. 5c), consistent with the peak of Gemin2ΔN39/SMN_Ge2BD_ alone (at ∼15.66 ml, Fig S3e). Comparing the staining intensity of 7 Sm proteins in lanes 13B-15B with that in lanes 11A-12B, we could estimate that the assembly of Sm core on U4-5’Δ was 2-fold more than on flU4-spacer-3’ss. This showed that Sm-core assembly is more efficient on the RNAs containing the snRNP code than on those containing the Sm site plus a linear 3’-sequence. This result is consistent with the previous report, in which a “des-stem” RNA (equivalent to the Sm site plus a 3’ single strand) was microinjected into the cytoplasm of Xenopus oocytes and its assembly efficiency into the Sm core was reduced by 2 folds (9).

The assembly efficiencies of the two types (types 1 and 2) of RNAs containing the snRNP code, with or without the 5’ adjacent SL of the Sm site, into the Sm core have little difference, as demonstrated by the incubation of equal molar amount of flU4 (type 1) and U4-5’Δ (type 2) with 7SΔN and D3(1-75)/B(1-91) followed by GFC (Fig. 5d). Similar amount of Sm cores were observed to assemble on flU4 and U4-5’Δ.

### Gemin2 serves as a checkpoint in Sm core assembly via negative cooperativity

Superposition of 7SΔN with U4 Sm core (40) (Fig. 6a, b) or U1 Sm core (39) (Data not shown) on SmF/E/G reveals that there is no clash of Gemin2’s N-terminal domain (NTD) with RNA, and on SmD1/D2 reveals that there is no clash of Gemin2’s CTD with RNA too. This indicates that Gemin2ΔN39 and RNA are not spatially exclusive. However, the binding of Gemin2ΔN39 on the periphery of 5Sm inhibits RNA assembly into the central RNA-binding pocket of 5Sm, allowing only the cognate snRNP code containing RNAs preferably to assemble into the Sm subcore. Moreover, the binding of cognate RNAs to 5Sm causes a “narrow-to-wide” conformational change of the latter, which decreases the binding affinity of Gemin2 to 5Sm and causes Gemin2 to dissociate from the Sm subcore. This phenomenon, in which Gemin2 and RNA are allosterically inhibitory on each other’s binding to 5Sm, is known as negative cooperativity. Although the concept of negative cooperativity has been well known, it has never been known as a mechanism in Sm core assembly. The widening of the conformation of 5Sm caused by cognate RNAs’ assembly also allows SmD3/B to join to further finish Sm core assembly. Therefore, in addition to the previously identified role of holding 5Sm, Gemin2 serves as a checkpoint in snRNP core assembly via a negative cooperativity mechanism, coupling RNA selection with Gemin2’s release, and ensuring that SmD3/B joins only after cognate RNAs’ assembly into Sm subcores.

**Figure 6.**
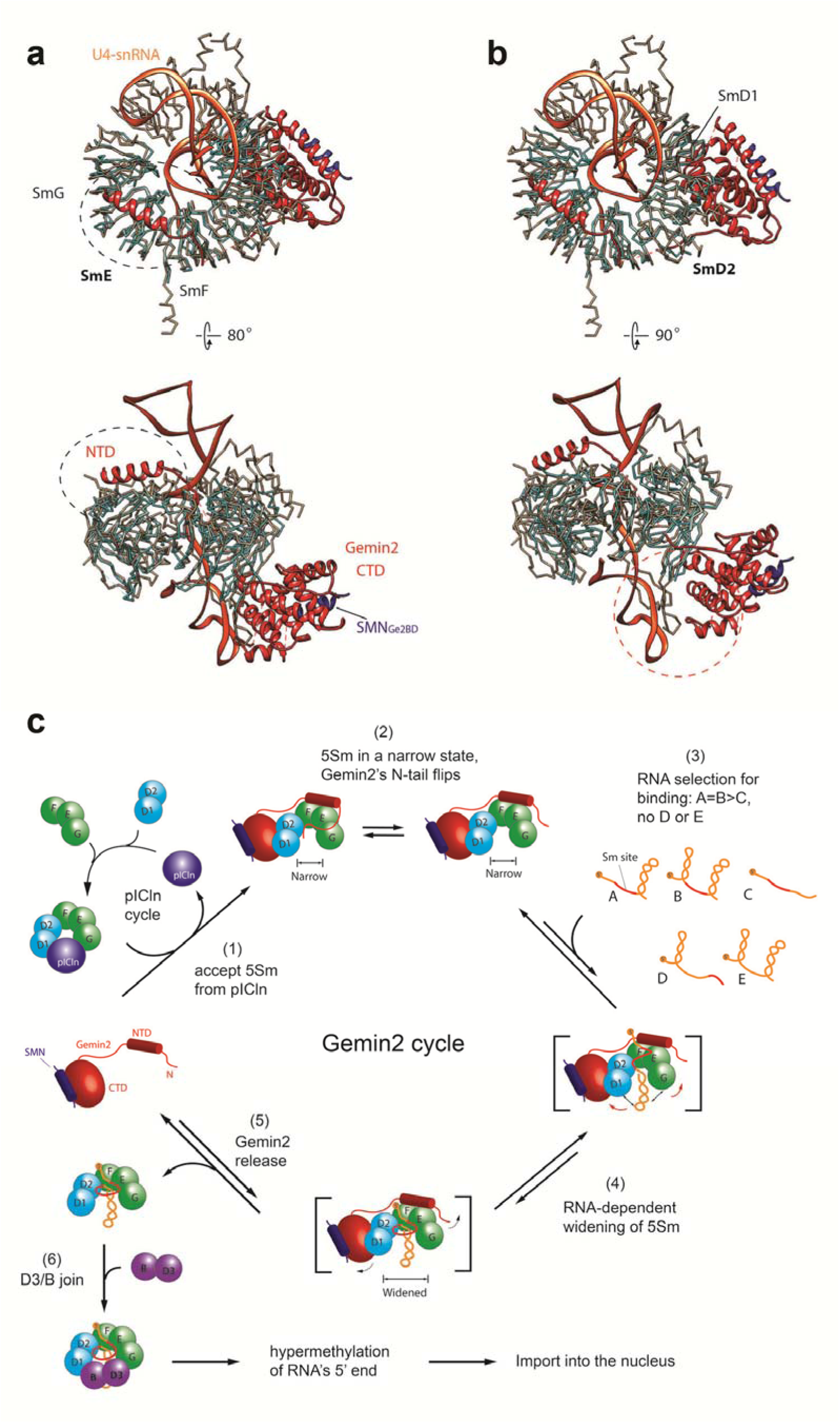
The bindings of Gemin2 and U4 snRNA to 5Sm are compatible sterically and the schematic model of Sm core assembly *in vivo*. Superposition of SmE (a) or SmD2 (b) of the 7SΔN complex with that of U4 snRNP core (PDB code 4WZJ)(40) reveals that Gemin2 is compatible with U4 snRNA in binding to 5Sm spatially, indicating allosteric, not competitive inhibition between Gemin2 and U4 snRNA in binding to 5Sm. The five Sm proteins in the 7SΔN complex are colored in cyan, and in the U4 snRNP core in grey. SMN_Ge2BD_ and Gemin2 are colored in blue and red respectively. U4 snRNA is colored in orange. (c) The mechanisms of Sm core assembly. The steps (2)-(6) are the new mechanisms discovered in this study. The step (1) is from literature (12, 15).

## DISCUSSION

In this study, we closely examined the assembly steps of the Sm core in the second phase, from the formation of SMN/Gemin2/5Sm, to the assembly of the Sm subcore, and finally to the completion of the Sm core by a combination of structural and biochemical approaches. We established that the narrow state of 5Sm bound by Gemin2/SMN is real and independent of Gemin2’s N-tail, and discovered its physiological role. We identified Gemin2’s second role in Sm core assembly in addition to being a holder of 5Sm—it serves as a checkpoint of the assembly by a negative cooperativity mechanism. By binding the outside of the horseshoe-shaped 5Sm and constraining it in a narrow conformation, Gemin2 helps 5Sm select RNA substrates, allowing preferably the cognate snRNAs, containing the snRNP code, to assemble into the inside of 5Sm to form subcores; the assembly of cognate RNAs widens 5Sm, which in turn causes Gemin2’s release and allows SmD3/B to join to complete the assembly of Sm cores. Non-cognate RNAs might be able to temporarily enter the central RNA-binding site of 5Sm, but are unable to widen 5Sm; the assembly stalls in the 7S intermediate state, which cannot go on to recruit SmD3/B and cause Gemin2’s release until the non-cognate RNAs inside 5Sm are replaced by cognate ones. Our proposed mechanism is schematically drawn in Figure 6c. This mechanism simultaneously provides answers to the two basic questions, how the SMN complex confers RNA assembly specificity, and how the SMN complex dissociates from the assembled Sm core. These findings cause a paradigm shift in our understanding of the mode of action of the SMN complex and snRNP assembly.

Since Sm core assembly *in vitro* is a spontaneous process, why eukaryotes evolve so many assembly chaperones in this process is a central question. Early studies have proved that these chaperones, especially the SMN complex, help the assembly in an highly specific way (8), and require RNA substrates containing both the nonameric Sm site and adjacent 3’-SL(9). People have been long probing the assembly specificity mechanism and found that pICln(12, 15), Gemin2’s N-tail(14) and Gemin5(29–34) are the candidates, however, which of the chaperones plays the major role to confer RNA assembly specificity and how it does have not yet been understood. pICln binds 5Sm in a closed ring to form the 6S complex, preventing premature and illicit assembly. However, it prevents any RNAs including the cognate ones to bind (12, 15). Similarly, although the N-tail of Gemin2, which we observed inside the RNA-binding pocket of 5Sm in our previous study, plays an inhibitory role in RNA-binding, it inhibits the binding of both correct and illicit RNAs (14). Therefore, both pICln and Gemin2’s N-tail are unable to serve as the specificity factor. Gemin5 of the SMN complex has long been considered as the specific factor and to play the role by direct binding to the snRNP code (29–34). This model is currently the dominating mechanism. Although Gemin5 is the first component of the SMN complex to bind precursor snRNAs and deliver them to the rest of the SMN complex in vertebrates, this model has several drawbacks. First, it cannot explain why snRNPs still assemble efficiently in many lower eukaryotes where no ortholog of Gemin5 is found (16, 35). Second, RNAi-mediated knockdown of Gemin5 has little effect on Sm core assembly (57, 58). Third, recent structural and biochemical studies of Gemin5-RNA interactions also provided evidence against this model: (1) only part of the Sm site, AUUU, much less than the snRNP code, is critical for RNA binding to Gemin5 (32–34); (2) Gemin5 can bind promiscuous RNAs, i.e., U1-tfs, which are the truncated U1 pre-snRNAs lacking the Sm site and the following SL (32). In contrast, our finding that Gemin2 achieves the high specificity of snRNP core assembly by a negative cooperativity mechanism can explain all these puzzles. First, Gemin2 is the most conserved component of the complex, from human to yeast (16, 59). Structure-guided sequence alignment of Gemin2 homologs in various species indicates that they all have the conserved F/E binding domain and D1/D2 binding domain, therefore would bind to 5Sm in the same way as does the human Gemin2 (14). Second, the minimal required RNA feature by the 7S complex is a Sm site plus a 3’-RNA, preferably a 3’-SL, matching the snRNP code identified previously (9), which is much more than that for binding to Gemin5. Third, our model can explain the previous *in vivo* and *in vitro* results. Using cell extracts and pull-down assays, the SMN complex was shown to assemble the major spliceosomal U-rich snRNPs, while total proteins (containing the 7 Sm proteins) also assembled other types of RNAs (8). The test condition was the purified SMN complex containing Sm proteins. With hindsight, we can readily deduce that the 5 Sm proteins were already bound by Gemin2, in the narrow, constrained state. In summary, it is the combination of 5Sm and Gemin2 rather than Gemin5 that mainly determines RNA specificity and recognizes the snRNP code. As Gemin5 is the first protein of the SMN complex in more complexed eukaryotes to bind snRNAs, it may play a role of preliminary screening of RNAs. It is interesting that under our *in vitro* experimental condition RNAs containing the snRNP code were merely about two-fold more efficient in Sm core assembly than RNAs just containing the Sm site in the middle (Fig. 5c). However, this efficiency is consistent with the previous *in vivo* experimental result, in which a radioactivity-labeled RNA containing the Sm site plus a 3’ single strand was observed to assemble into the Sm core by 50% efficiency after microinjected into the cytoplasm of Xenopus oocytes and pulled-down by anti-Sm antibody (9). This consistency further supports that our findings reflect the *in vivo* situation. As for how to understand that the SMN complex dominantly assemble the Sm core on snRNAs instead of non-cognate RNAs inside cells, there may be two additional contributing factors: (1) high abundance of snRNAs in cells compared with other non-cognate RNAs which containing just the Sm site (60, 61); (2) in other non-cognate RNAs containing only the Sm site, the single strand 3’ to the Sm site might be a binding site of other proteins, which would block the Sm core assembly.

The mechanism of the SMN complex’s dissociation from the Sm core has long been elusive. Our findings provide the first model on it. Because Gemin2 is the major protein to bind Sm proteins (14) and therefore, we reason that how the SMN complex is released depends mostly on how Gemin2 is released. We had hypothesized that Gemin2 were released by a cleavage at the connecting loop between its NTD and CTD inside cells. But we found that Gemin2 is entirely intact in Hela cells when we tagged Gemin2 at both its N- and C-termini in Hela cells and checked its tags by Western blot (Data not shown). How Gemin2 can dissociate intact from the Sm core had puzzled us for a while. In this study, we found that Gemin2 is partially released upon cognate RNAs’ assembly into 5Sm and completely upon Sm-core formation. Accordingly, the SMN complex should stay in the cytoplasm if Gemin2’s detachment is the only mechanism for the entire SMN complex’s release. In Hela cells, most of the SMN complex components, including Gemin2, were observed also in the nucleus, and concentrated in CBs (29). How to explain this apparent disparity? First of all, we notice that the concentrations of most of the SMN complex proteins are higher in the cytoplasm than in the nucleus (29). This tells that most of the SMN complex may dissociate from the Sm core and recycle in the cytoplasm, which is consistent with our proposed model. In addition, we also know that the SMN protein contains a tudor domain, which is able to bind the methylated RG-rich tails of SmD1, D3 and B (62). It is likely that small portion of the SMN complex remain tethered to Sm cores through these interactions and follows Sm cores into the nucleus. In the CBs, the methylated arginine residues in the CB-hallmark protein coilin can bind SMN (63). It is possible that coilin competes with the methylated RG-rich tails of SmD1, D3 and B, releasing Sm cores from the SMN complex completely while keeping the SMN complex in CBs.

Because of the highest conservation of Gemin2 among the SMN complex in eukaryotes (16, 59), we propose that this negative allosteric mechanism mediated by Gemin2 is a fundamental mechanism of Sm-core assembly in all eukaryotes. The fact that only orthologue of Gemin2, Brr1, is found to play an important role in Sm-core assembly in one of the simplest eukaryote, *S. cerevisiae* supports this idea (16, 64). Recent genetic study in *S. cerevisiae* provided details of interactions of Brr1 with SmD1, SmD2, SmF, and SmE of the Sm core, but not with SmG, SmB and SmD3 (65), further supporting the similarity between Brr1 and Gemin2 in Sm-core assembly. In this simple eukaryote, the Gemin2 cycle mechanism might explain all the mechanism. This finding also provides further clues for the evolution of the Sm-core assembly chaperone system. This work would facilitate further mechanistic study of other components of the SMN complex in snRNP assembly in high eukaryotes. For example, the structure of Gemin6/7 resembles a Sm heterodimer and it has been suggested that Gemin6/7 serve as a surrogate to bind to 5Sm in the position of SmD3/B (66). Our study makes this suggestion less likely probable because the narrow 5Sm disables any Sm-fold dimer to join, and once a cognate RNA binds to 5Sm and widens 5Sm, SmD3/B can readily join to finish the assembly efficiently. There is no reason for Gemin6/7 to bind first. So Gemin6/7 may play a different role, which awaits further investigation.

In addition, we predict that this mechanism likewise apply to the assembly of U7 snRNP core, which has a variation of Sm protein components and Sm site but is assembled by the SMN complex and requires a 3’-SL adjacent to the Sm site (43–45).

Furthermore, our finding also facilitates pathogenesis study of SMA and development of therapeutics. The demonstration of Gemin2 offering the basic mechanism of Sm core assembly provides the possibility to develop strategies to assemble Sm cores without the SMN protein (For example, using the Brr1 protein mutant which is enabled to bind human 5Sm). These strategies could be used to test if SMA is totally caused by deficiency of Sm core assembly and to develop possible therapeutics targeted on Sm core assembly.

The major method used in this study to characterize protein-RNA interaction is GFC followed by SDS-PAGE, a two-dimensional separation approach. It is an approach commonly used in protein-protein interactions, yet much rarely used in protein-RNA interaction studies compared with EMSA. However, it is superior to one-dimensional EMSA when multiple-component systems are studied, as illustrated here by discovering the release of Gemin2 from Sm subcores, which escaped from many previous studies by EMSA (8,12,15). So this two-dimensional approach is generally applicable for other protein-RNA and protein-DNA interactions, especially studying nucleotides interaction with multiple-protein components. However, like in our present study, applying this approach has to meet the caveats: (1) requirement of the tested complexes being substantially stable, (2) requirement of much more amount of purified proteins and nucleic acids, and (3) requirement of more sophisticated fast protein liquid chromatography (FPLC) system equipped with multiple-wavelength detector.

In summary, we discovered a new mechanism in the assembly of snRNP core—the negative cooperativity between Gemin2 and RNA in binding to 5Sm, which simultaneously answers two basic questions in snRNP assembly: how does the SMN complex achieves high RNA assembly specificity and how it is released? This finding provides better understanding of the mode of action of the SMN complex and the snRNP assembly, including all spliceosomal snRNPs and likely U7 snRNP as well.

## ACCESSION CODES

Atomic coordinates and structural factors for the reported crystal structures of complexes A, B and C have been deposited with the Protein Data Bank under accession codes 5XJQ, 5XJS and 5XJR.

## SUPPLEMENTARY DATA

Supplementary Data are available online.

## ACKNOWLEDGEMENTS

We thank Drs. Gideon Dreyfuss, Klaus Hartmuth, Reinhard Luhrmann and Ruben J. Cauchi for reading the manuscript and providing comments. We thank the staff of the beamlines BL17U1 and BL19U1 at the National Facility for Protein Science (NFPS) and Shanghai Synchrotron Radiation Facility, Shanghai, People’s Republic of China, for assistance during crystal diffraction data collection.

## AUTHOR CONTRIBUTIONS

H. Yi crystallized the complexes. H. Yi and L. Mu performed the biochemical assays. C. Shen, X. Kong, Y. Wang and Y. Hou participated in the project. R. Zhang conceived, designed and supervised the project, solved the crystal structures and wrote the paper.

