## supplemental materials for "Negative Cooperativity between Gemin2 and RNA provides Insights into RNA Selection and the SMN Complex’s Release in snRNP Assembly"

Containing

7 supplemental tables (Tables S1-7),

8 supplemental figures (Figures S1-8)

**Table S1. Data Collection and Refinement Statistics**

| **Variable protein components*** | | | |
|---|---|---|---|
| | Complex A | Complex B | Complex C |
| Gemin2 | FL(1-280) | ΔN39 (40-280) | ΔN39 (40-280) |
| SmD1 | ΔC(1-82) | ΔC(1-82) | ΔC(1-82) |
| SmG | Yes | No | Yes |
| **Data collection** | | | |
| Wavelength (Å) | 0.97853 | 0.97853 | 0.97853 |
| Space group | $P2_12_12_1$ | $P2_12_12_1$ | $P2_12_12_1$ |
| Unit cell (Å): | 83.32, | 83.17, | 82.96, |
| a, b, c | 115.76, | 114.09, | 114.12, |
| | 128.21 | 125.39 | 130.14 |
| Highest resolution (Å)[$]: | | | |
| a*, b*, and c* | 3.3, 3.5, 3.9 | 3.4, 3.4, 4.0 | 3.1, 3.2, 3.6 |
| Unique reflections | 15470 | 14569 | 18768 |
| Completeness (%) | 98.2 (94.6)[a] | 99.5(97.9)[b] | 97.8(95.2)[c] |
| | 79.2(15.9)[d] | 84.2(35.5)[e] | 83.1(28.8)[f] |
| $R_{meas}$ | 0.154 (0.905) | 0.233(0.914) | 0.166(0.885) |
| $R_{pim}$ | 0.048 (0.261) | 0.087(0.297) | 0.056(0.243) |
| Mean I/σ | 13.6 (2.2) | 12.9(2.0) | 10.4(2.0) |
| Redundancy | 12.4 (12.3) | 10.8 (8.7) | 12.4(12.7) |
| **Refinement Statistics** | | | |
| Resolution range(Å) | 48-3.3 | 47-3.4 | 47-3.1 |
| R factor (%) | 20.5 | 19.1 | 17.2 |
| $R_{free}$ factor (%) | 26.5 | 24.5 | 24.0 |
| Number of reflections | 15470 | 14569 | 18768 |
| Number of atoms | 4793 | 4346 | 4890 |
| Rmsd bond length (Å) | 0.0112 | 0.0122 | 0.0129 |
| Rmsd bond angles (°) | 1.576 | 1.624 | 1.477 |
| Ramachandran plot (%)[#]: | | | |
| Favored, | 93.9, | 91.6, | 94.7, |
| additional allowed, | 5.9, | 7.5, | 4.6, |
| disallowed | 0.2 | 0.9 | 0.7 |
| PDB code | 5XJQ | 5XJS | 5XJR |

* Complexes A-C all contain SMN(residues 26-62), SmD2, SmF and SmE.

[$] Ellipsoidal truncation was performed on each data set due to serious anisotropic diffraction.

[a-c] For resolution range of 48-3.9, 47-4.0 and 47-3.6 Å respectively. The number in the parenthesis corresponds to the highest-resolution shell of 4.04-3.90, 4.12-4.00 and 3.72-3.60 Å respectively.

[d-f] For resolution range of 48-3.3, 47-3.4 and 47-3.1 Å respectively. The number in the parenthesis corresponds to the highest-resolution shell of 3.4-3.3, 3.52-3.40 and 3.2-3.1 Å respectively.

[#] gained from Coot program.

**Table S2. RNA sequences used in this study.**

| Name | RNA sequence |
|---|---|
| 9nt | AAUUUUUGA |
| 3’Sm | GGGAAUUGAAAACUUUUCCCAAUACCCCAAUUUUUGA |
| U4 | GGGAAUUGAAAACUUUUCCCAAUACCCCGCCGUGACGACUUGCAAUAU<br>AGUCGGCACUGGCAAUUUUUGACAGUCUCUACGGAGACUG |
| U4ΔSm | GGGAAUUGAAAACUUUUCCCAAUACCCCGCCGUGACGACUUGCAAUAU<br>AGTCGGCACUGGCAACCCCCGACAGUCUCUACGGAGACUG |
| U4-3’ss | GGGAAUUGAAAACUUUUCCCAAUACCCCGCCGUGACGACUUGCAAUAU<br>AGUCGGCACUGGCAAUUUUUGACAGUCUCUACGCUCUGAC |
| U4-3’Δ | GGGAAUUGAAAACUUUUCCCAAUACCCCGCCGUGACGACUUGCAAUAU<br>AGUCGGCACUGGCAAUUUUUGAC |
| U4-5’ss | GGGAAUUGAAAACUUUUCCCAAUACCCCAAUUUUUGACAGUCUCUACG<br>GAGACUG |
| U4-5’Δ | GGCAAUUUUUGACAGUCUCUACGGAGACUG |
| U4-5’Δ-3’ss | GGCAAUUUUUGACAGUCUCUACGCUCUGAC |
| flU4 | GGGCAGCUUUGCGCAGUGGCAGUAUCGUAGCCAAUGAGGUCUAUCCGA<br>GGCGCGAUUAUUGCUAAUUGAAAACUUUUCCCAAUACCCCGCCGUGAC<br>GACUUGCAAUAUAGUCGGCACUGGCAAUUUUUGACAGUCUCUACGGAG<br>ACUG |
| flU4ΔSm | GGGCAGCUUUGCGCAGUGGCAGUAUCGUAGCCAAUGAGGUCUAUCCGA<br>GGCGCGAUUAUUGCUAAUUGAAAACUUUUCCCAAUACCCCGCCGUGAC<br>GACUUGCAAUAUAGUCGGCACUGGCAACCCCCGACAGUCUCUACGGAG<br>ACUG |
| flU4-spacer | GGGCAGCUUUGCGCAGUGGCAGUAUCGUAGCCAAUGAGGUCUAUCCGA<br>GGCGCGAUUAUUGCUAAUUGAAAACUUUUCCCAAUACCCCGCCGUGAC<br>GACUUGCAAUAUAGUCGGCACU<u>CCG</u>AAUUUUUGACAGUCUCUACGGAG<br>ACUG |
| flU4-spacer-3’ss | GGGCAGCUUUGCGCAGUGGCAGUAUCGUAGCCAAUGAGGUCUAUCCGA<br>GGCGCGAUUAUUGCUAAUUGAAAACUUUUCCCAAUACCCCGCCGUGAC<br>GACUUGCAAUAUAGUCGGCACU<u>CCG</u>AAUUUUUGACAGUCUCUACGCUC<br>UGAC |

Red, Sm site. Blue, ΔSm. <u>Cyan, spacer</u>.

**Table S3. The components and amounts used for gel filtration chromatographic (GFC) assay.**

| Input components (RNA colored in blue) | Amount used (molar ratio in last brackets) |
|---|---|
| D1s/D2 + F/E/G (5Sm) | 200μg + 200μg (1:1) |
| Gemin2ΔN39 | 200μg |
| D1s/D2 | 1mg |
| F/E/G | 1mg |
| 3'Sm | 10μg |
| 5Sm + 3'Sm | 400μg + 100μg (1:1) |
| 5Sm + 9nt | 400μg + 120μg (about 1:5) |
| Gemin2ΔN39/$SMN_{Ge2BD}$ + D1s/D2 + F/E/G (7SΔN preparation) | 0.6mg + 1mg + 1mg (1:2:2) |
| 7SΔN + 3'Sm | 160μg + 40μg (about 1:1.5) |
| 7SΔN + U4 | 160μg + 60μg (1:1) |
| (7SΔN + D3/B) + U4 | (160μg + 60μg) + 60μg (1:1.5:1) |
| 7SΔN + U4ΔSm | 160μg + 60μg (1:1) |
| 7SΔN + U4-3'Δ | 160μg + 50μg (1:1) |
| 7SΔN + U4-5'Δ | 160μg + 40μg (1:2) |
| U4-5'Δ | 5μg |
| 7SΔN + U4-3'ss | 160μg + 60μg (1:1) |
| 7SΔN + U4-5'ss | 160μg + 40μg (1:1) |
| 7SΔN + U4-5'Δ-3'ss | 80μg + 40μg (1:4) |
| 7SΔN + U4-5'Δ-3'ss + U4-5'Δ | 80μg + 15μg + 15μg (1:1.5:1.5) |
| 5Sm + U4-5'Δ | 400μg + 40μg (2:1) |
| 5Sm + U4-5'ss | 400μg + 80μg (2:1) |
| flU4 | 10μg |
| 7SΔN + flU4 | 120μg + 120μg (1:1.6) |
| 7SΔN + flU4ΔSm | 100μg + 100μg (1:1.6) |
| 7SΔN + flU4-spacer | 120μg + 120μg (1:1.6) |
| 7SΔN + flU4-spacer-3'ss | 80μg + 80μg (1:1.6) |
| (7SΔN + D3/B) + flU4 | (120μg + 60μg) + 120μg（1:1.5:1.6） |
| (7SΔN + D3/B) + flU4-spacer-3'ss | (80μg + 40μg) + 80μg (1:2:1.6) |
| 7SΔN + (flU4-spacer + U4-5'Δ-3'ss) | 80μg + (80μg + 16μg) (1:1.6:1.6) |
| 7SΔN + (flU4-spacer-3'ss + U4-5'Δ) | 80μg + (80μg + 16μg) (1:1.6:1.6) |
| (7SΔN + D3/B) + (flU4-spacer-3'ss + U4-5'Δ) | (80μg + 40μg) + (80μg + 16μg) (1:2:1.6:1.6) |
| (7SΔN + D3/B) + (flU4 + U4-5'Δ) | (100μg + 50μg) + (100μg + 20μg) (1:2:1.6:1.6) |

**Table S4. The GFC elution positions of the RNAs, proteins, and their complexes studied.**

| Components | Elution position (ml) | +7SΔN: Elution position (ml) | +5Sm: Elution position (ml) | +7Sm: Elution position (ml) |
|---|---|---|---|---|
| 9nt | ~19.6 | Not formed | 14.37 | Not tested |
| 3'Sm | 16.64 | Not formed | 13.51 | Not tested |
| U4 | 14.75 | 13.31 | Not tested | 13.43 |
| U4ΔSm | 14.91 | Not formed | Not tested | Not tested |
| U4-3'ss | 14.48 | 13.11 | Not tested | Not tested |
| U4-3'Δ | 15.66 | Not formed | Not tested | Not tested |
| U4-5'ss | 15.67 | 13.65(RNA+5Sm) | 13.50 | Not tested |
| U4-5'Δ | 17.48 | 14.29(RNA+5Sm) | 14.27 | Not tested |
| U4-5'Δ-3'ss | 16.62 | 13.97 | Not tested | Not tested |
| flU4 | 13.08 | 12.26 | Not tested | 12.31 |
| flU4ΔSm | 12.93 | Not formed | Not tested | Not tested |
| flU4-spacer | 12.97 | 12.24(RNA+5Sm) | Not tested | Not tested |
| flU4-spacer-3'ss | 12.90 | ~12.11 | Not tested | 12.14 |
| 7SΔN | 13.78 | ---------- | ---------- | ---------- |
| 7S | 13.61 | ---------- | ---------- | ---------- |
| D1s/D2 | 16.63 | ---------- | ---------- | ---------- |
| F/E/G | 14.29(dimer), 16.62(monomer) | ---------- | ---------- | ---------- |
| D1s/D2/F/E/G(5Sm) | 15.06 | ---------- | ---------- | ---------- |
| Gemin2ΔN39/$SMN_{Ge2BD}$ | 15.66 | ---------- | ---------- | ---------- |

5Sm indicates D1s/D2 and F/E/G. 7Sm indicates 5Sm and D3(1-75)/B(1-91).

**Table S5. The molar extinction coefficients of the proteins, protein complexes and RNAs studied.**

| Sample | Length | MW(Da) | Molar Extinction Coefficient (mol/L·cm$^{-1}$) | |
|---|---|---|---|---|
| | | | at 280 nm | at 260 nm |
| **Protein:** | | | | |
| SmD1s(1-82) | 82 | 9287.57 | 1280 | 640 |
| SmD2 | 119 | 13583.36 | 7210 | 3605 |
| SmF | 86 | 9724.38 | 12210 | 6105 |
| SmE | 92 | 10802.89 | 10930 | 5465 |
| SmG | 77 | 8552.85 | 120 | 60 |
| SmD3(1-75) | 75 | 8465.17 | 4080 | 2040 |
| SmB(1-92) | 92 | 10508.92 | 1640 | 820 |
| D1s/D2 | 201 | 22870.93 | 8490 | 4245 |
| SmF/E/G | 255 | 29080.12 | 23260 | 11630 |
| D3/B | 167 | 18974.09 | 5720 | 2860 |
| 5Sm | 657 | 74821.98 | 31750 | 15875 |
| 7Sm | 824 | 93796.07 | 37470 | 18735 |
| Gemin2ΔN39 | 242 | 27347.49 | 36970 | 18485 |
| SMN(26-62) | 37 | 4042.12 | 7090 | 3545 |
| Gemin2ΔN39/SMN(26-62) | 279 | 31389.61 | 44060 | 22030 |
| 7SΔN | 936 | 106211.59 | 75810 | 37905 |
| **RNA:** | | | | |
| 9nt | 9 | 3043 | 40909 | 81818 |
| 3'Sm | 37 | 12017.5 | 168182 | 336364 |
| U4 | 88 | 28363 | 400000 | 800000 |
| U4-3'ss | 88 | 28363 | 400000 | 800000 |
| U4-3'Δ | 71 | 22914.5 | 322727.5 | 645455 |
| U4-5'ss | 55 | 17786.5 | 250000 | 500000 |
| U4-5'Δ | 30 | 9774 | 136363.5 | 272727 |
| flU4 | 148 | 47593 | 672727 | 1345454 |
| flU4ΔSm | 148 | 47593 | 672727 | 1345454 |
| flU4-spacer | 148 | 47593 | 672727 | 1345454 |
| flU4-spacer-3'ss | 148 | 47593 | 672727 | 1345454 |

Note: The molar extinction coefficient (MEC) at 280nm of protein is calculated by Vector NTI software (Thermo Fisher Scientific). The MEC at 260nm of each protein is estimated as the half of the MEC at 280nm. The MEC of protein complex (in red) is simply the sum of its components.

The MEC at 260nm of RNA (in blue) is calculated on the basis of 1 $OD_{260}$ ssRNA=40μg/ml=0.11mM (in nucleotides) and its length. The equation is MEC (at 260nm) = 1000 X RNA length/0.11. The MEC at 280nm of RNA is estimated as the half of the MEC at 260nm.

**Table S6. The calculated concentrations of RNAs and RNA-protein complexes from GFC for single RNA's assembly into Sm subcore and core.**

| the sample incubated with 7SΔN for GFC | OD(mAu) at 260nm | | | MEC [at 260nm] | | | Concentration(nM) | | |
|---|---|---|---|---|---|---|---|---|---|
| | RNA-protein | RNA | 7SΔN* | RNA-protein# | RNA | 7SΔN | RNA-protein | RNA | 7SΔN* |
| U4 | 24.3 | 85.1 | 9.6 | 837905 | 800000 | 37905 | 29 | 106 | 253 |
| U4ΔSm | 0.0 | 74.4 | 9.6 | 837905 | 800000 | 37905 | 0 | 93 | 253 |
| U4-3'ss | 47.9 | 68.4 | 10.5 | 837905 | 800000 | 37905 | 57 | 85 | 278 |
| U4-3'Δ | 0.0 | 51.1 | 9.6 | 683360 | 645455 | 37905 | 0 | 79 | 253 |
| U4-5'ss | 37.9 | 79.8 | 11.0 | 515875 | 500000 | 37905 | 73 | 160 | 289 |
| U4-5'Δ | 32.3 | 29.0 | 14.3 | 288602 | 272727 | 37905 | 112 | 106 | 376 |
| flU4 | 101.1 | 228.0 | 11.0 | 1383359 | 1345454 | 37905 | 73 | 169 | 289 |
| flU4ΔSm | 0.0 | 174.5 | 17.1 | 1383359 | 1345454 | 37905 | 0 | 130 | 452 |
| flU4-spacer | 203.0 | 311.7 | 8.2 | 1361329 | 1345454 | 37905 | 149 | 232 | 217 |
| flU4-spacer-3'ss | 81.2 | 119.7 | 2.9 | 1361329 | 1345454 | 37905 | 60 | 89 | 77 |
| flU4+D3/B | 248.3 | 145.1 | 8.0 | 1364189 | 1345454 | 37905 | 182 | 108 | 210 |
| flU4-spacer-3'ss + D3/B | 120.2 | 112.2 | 2.9 | 1364189 | 1345454 | 37905 | 88 | 83 | 77 |

*Due to the lower MEC of 7SΔN than those of RNAs, the OD values of 7SΔN, which are at the similar level to the background, is unable to be estimated correctly compared with those of RNAs and RNA-protein complexes.

### The MEC of RNA-protein is calculated by addition of the MECs of RNA and 7SΔN (or 5Sm, or 7Sm. Due to the dominancy of RNA in MEC of RNA-protein complex, using any MEC of the three protein complexes gives rise to almost identical calculated concentrations for RNA-protein complexes.)

**Table S7. The calculated concentrations of RNAs and RNA-protein complexes from GFC for the competition of two RNAs' assembly into Sm subcores and cores.**

| the sample incubated with 7SΔN for GFC | OD(mAu) at 260nm | | | | MEC [at 260nm] | | | | Concentration (nM) | | | |
|---|---|---|---|---|---|---|---|---|---|---|---|---|
| | large RNA-protein | Large RNA | short RNA-protein* | short RNA* | large RNA-protein | Large RNA | short RNA-protein | short RNA | large RNA-protein | Large RNA | short RNA-protein* | short RNA* |
| flU4-spacer + U4-5'Δ-3'ss | 122.3 | 141.6 | 26.0 | 29.9 | 1361329 | 1345454 | 288602 | 272727 | 90 | 105 | 90 | 110 |
| flU4-spacer-3'ss + U4-5'Δ | 78.4 | 123.8 | 30.3 | 27.9 | 1361329 | 1345454 | 288602 | 272727 | 58 | 92 | 105 | 102 |
| flU4-spacer-3'ss + U4-5'Δ + D3/B | 75.8 | 69.1 | 43.3 | 21.9 | 1364189 | 1345454 | 288602 | 272727 | 56 | 51 | 150 | 80 |
| flU4 +U4-5'Δ + D3/B | 159.2 | 154.3 | 21.6 | 19.9 | 1364189 | 1345454 | 288602 | 272727 | 117 | 115 | 75 | 73 |

*The OD values of short RNAs and short RNA-protein complexes, which are at the similar level to the background, are unable to be estimated correctly compared with those of large RNAs and large RNA-protein complexes.

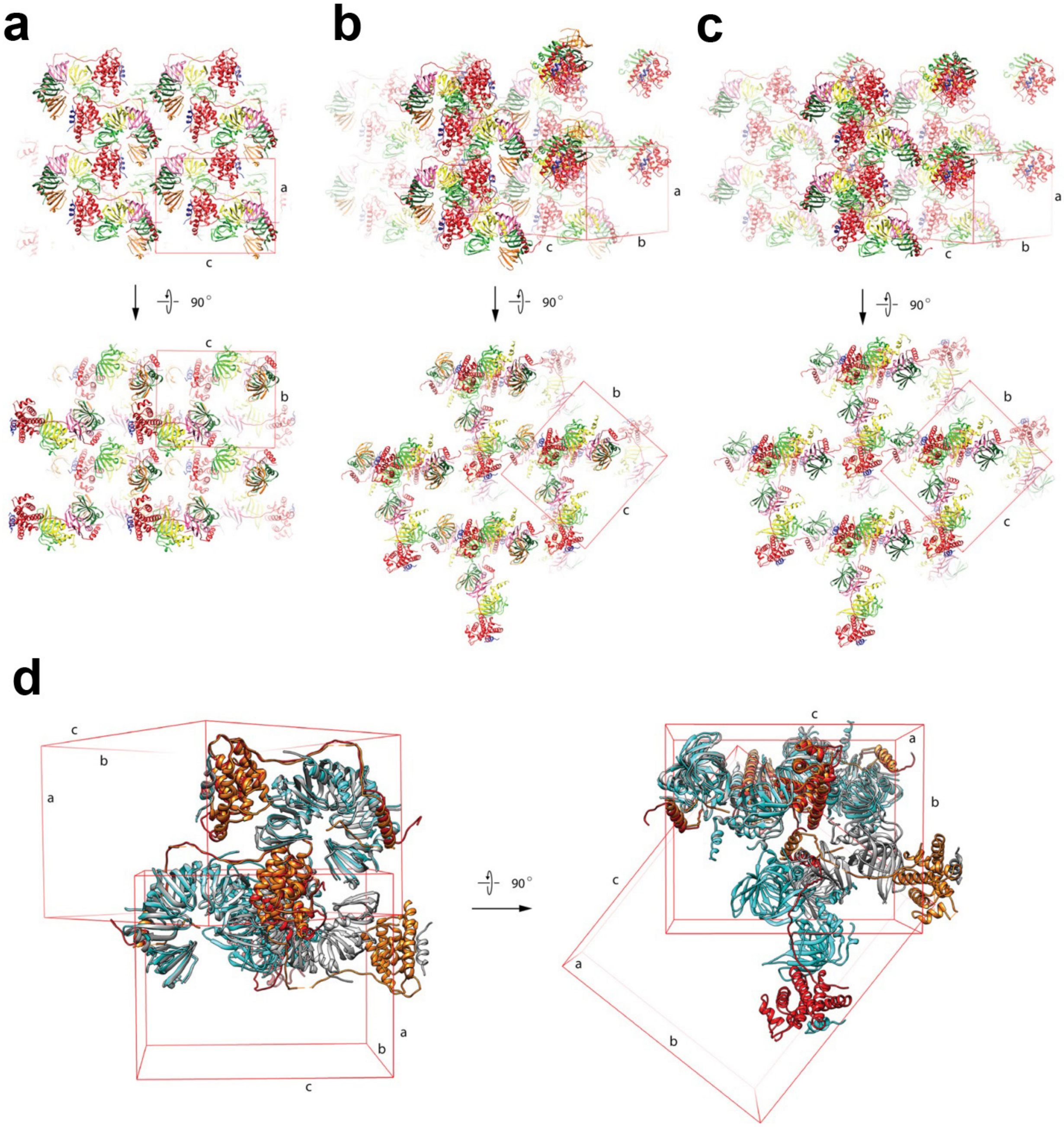

**Fig. S1. Crystal packing comparison of three complexes.** Two views of the crystal packing for each of 3 complexes: (a) the 7S complex from the previous study (3S6N), (b) Complex A and (c) Complex B. The five Sm proteins, D1, D2, F, E and G are colored in green, lemon, pink, dark green and orange respectively. Gemin2 and $SMN_{Ge2BD}$ are colored in red and blue respectively. Unit cells and axis are showed. (d) Comparison of 3S6N (5Sm in light gray and Gemin2 in orange) with Complex A (5Sm in cyan and Gemin2 in red) in crystal packing. Two molecules of the complexes are superimposed in direction a.

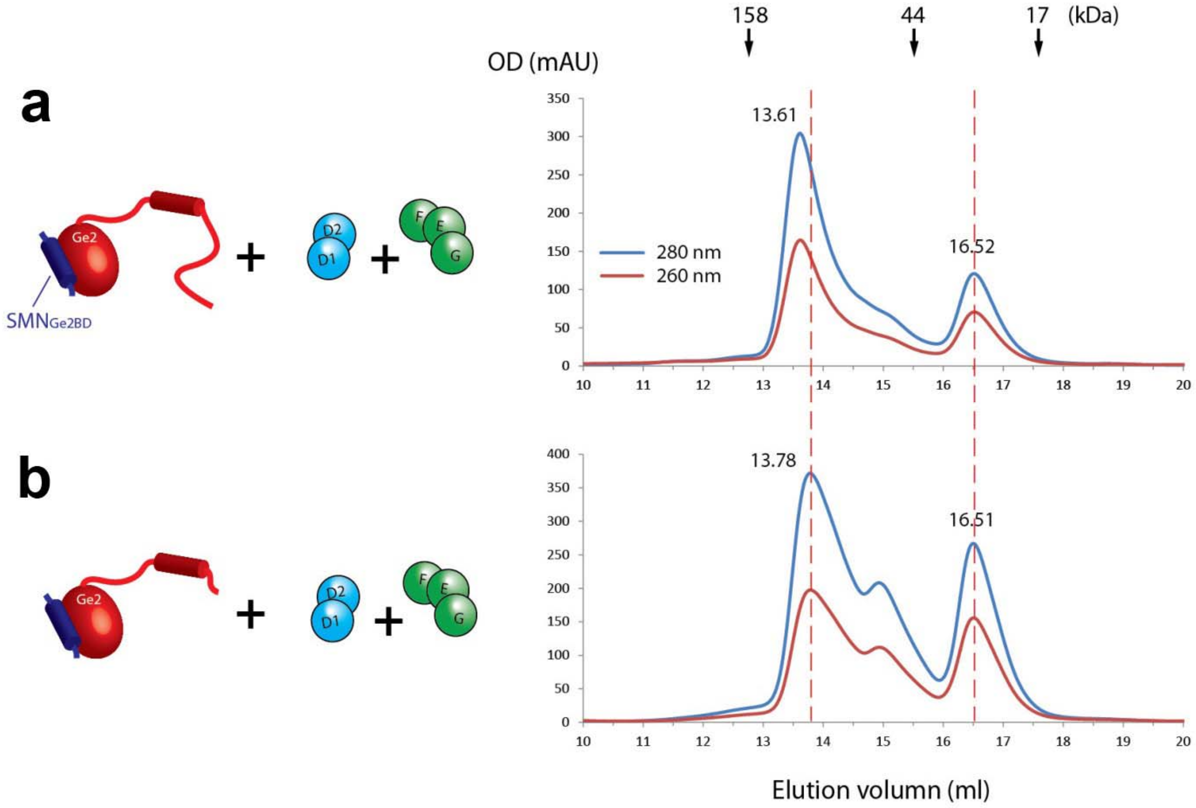

**Fig. S2. The gel filtration profiles of 7S and 7SΔN.** Reconstitution of 7S or 7SΔN was made by mixing extra amount of the 2 hetero-oligomeric Sm subcomplexes, D1s/D2 and F/E/G, with (a) Gemin2/SMN$_{Ge2BD}$ or (b) Gemin2ΔN39/SMN$_{Ge2BD}$ respectively followed by gel filtration chromatography separation (right panels). For each complex, one representative result from three independent experiments is shown. The input components are showed in cartoon (left panels). The positions of standard proteins are indicated at the top. Peak positions (ml) are indicated. The extra amount of D1s/D2 and F/E/G serve as inner controls (eluted at 16.5 ml). They also form small amount of D1s/D2/F/E/G heteropentamer (5Sm) at about 15.0 ml. The peaks of 7S and 7SΔN appeared at 13.61 and 13.78 ml respectively.

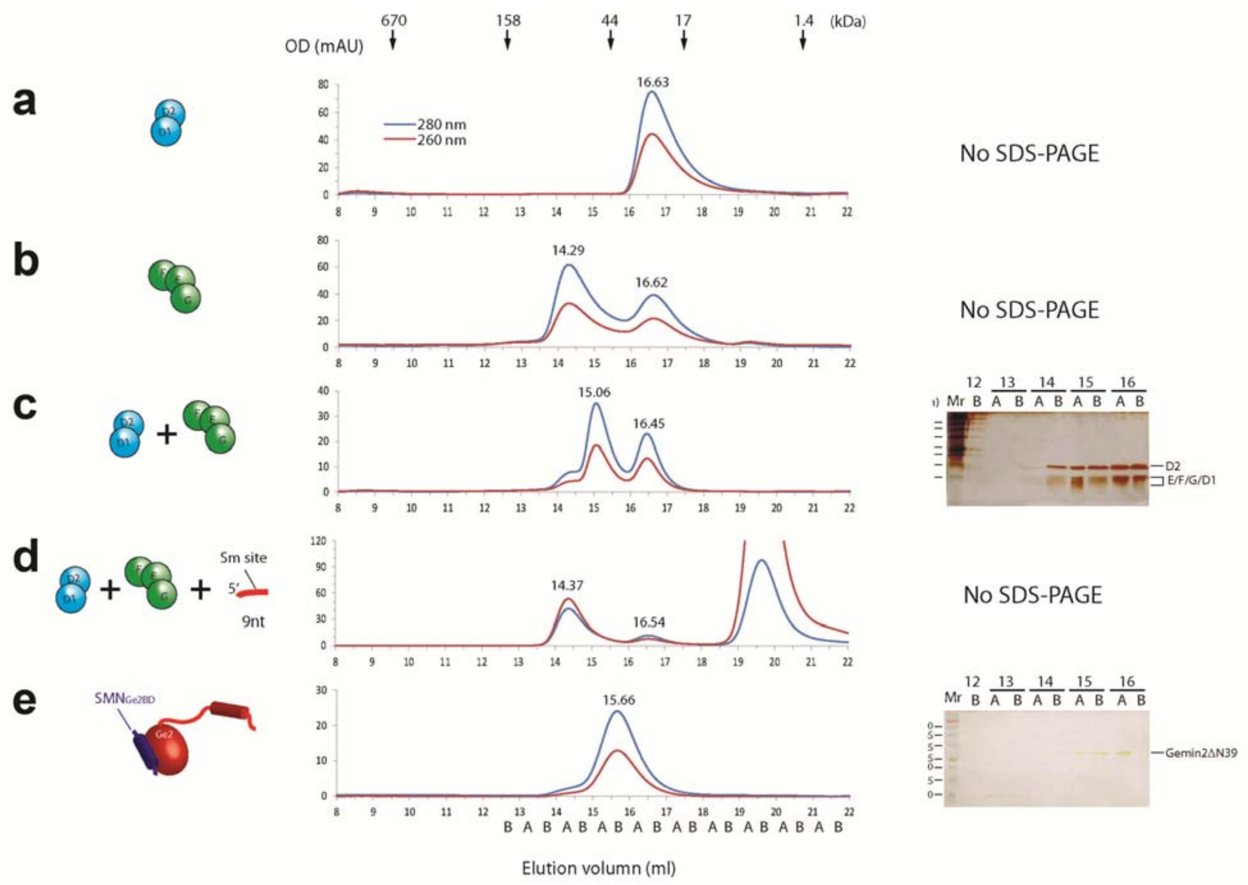

**Fig. S3. The gel filtration profiles of different complexes.** Different complexes (in cartoon, left panels), SmD1s/D2 (a), SmF/E/G (b), mixture of SmD1/D2 and SmF/E/G (c), mixture of 9nt and the 5 Sm proteins (d), or SMN(26-62)/Gemin2ΔN39 (e), was separated by gel filtration chromatography (middle panels) and individual fractions were analyzed by SDS-PAGE and silver staining (right panels). For each, one representative result from at least two independent experiments is shown. The positions of standard proteins are indicated at the top. Fractions (A &B) are named on the basis of volume positions (bottom). SmF/E/G (e) is the same as in Figure 2 and showed here for easy comparison. The SMN(26-62) fragment is too small to be visible on the SDS-PAGE in panel (e).

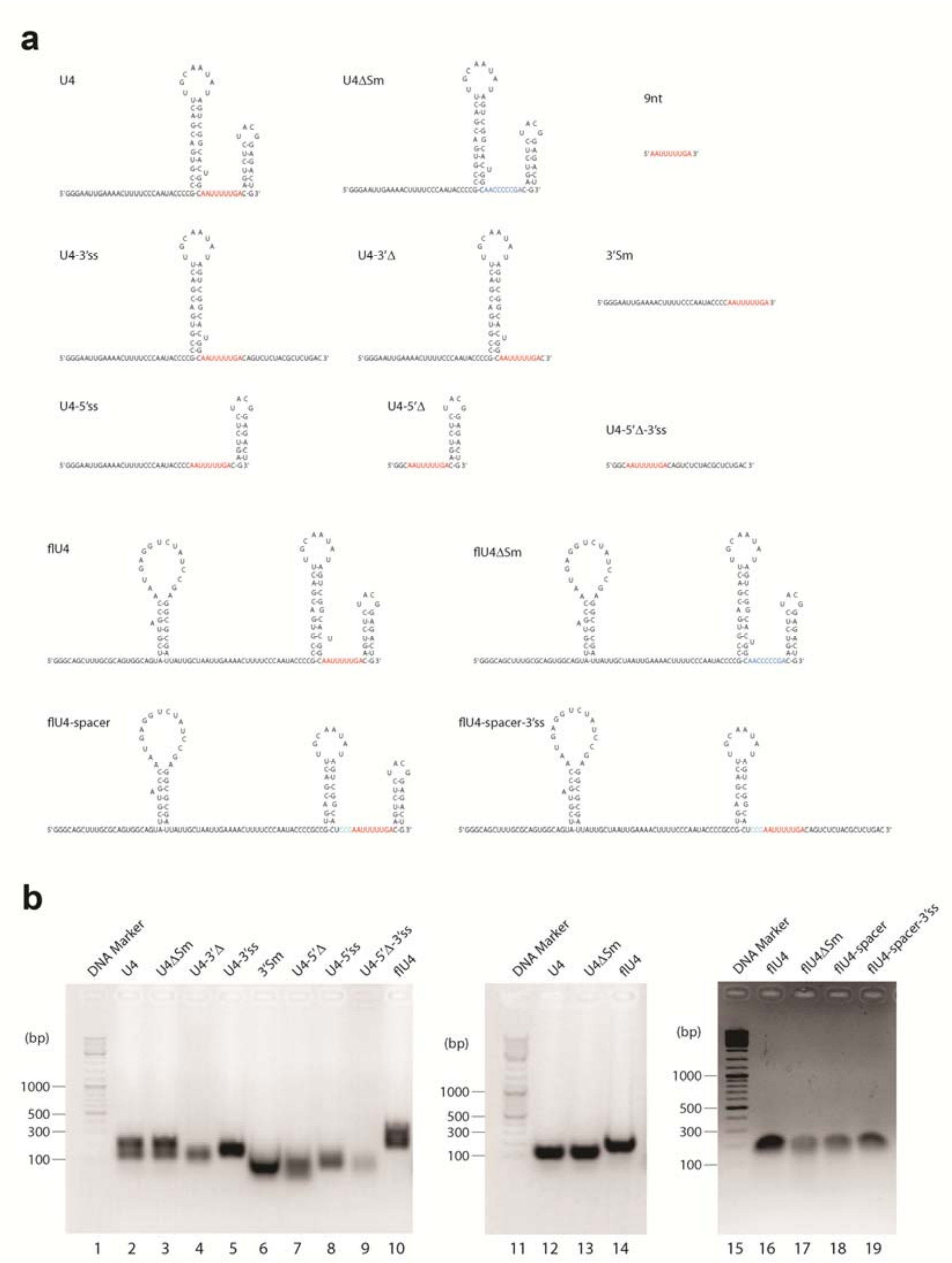

**Fig. S4. The RNAs used in this study.** (a) The sequence and secondary structures of the RNAs. Sm site is colored in red, ΔSm in blue, and the 3nt spacer in cyan (same as in cartoon). (b) The agarose gel electrophoresis of the *in vitro* transcribed RNAs. The RNAs in lanes 2-10 were not specially treated before loading on the gel. The three RNAs, U4, U4ΔSm and flU4, which were double-bands in lanes 2, 3 and 10, and flU4 and its three derivatives, were incubated at 65°C for 10 min and cooled to room temperature before loading on the gel (lanes 12-14 and 16-19).

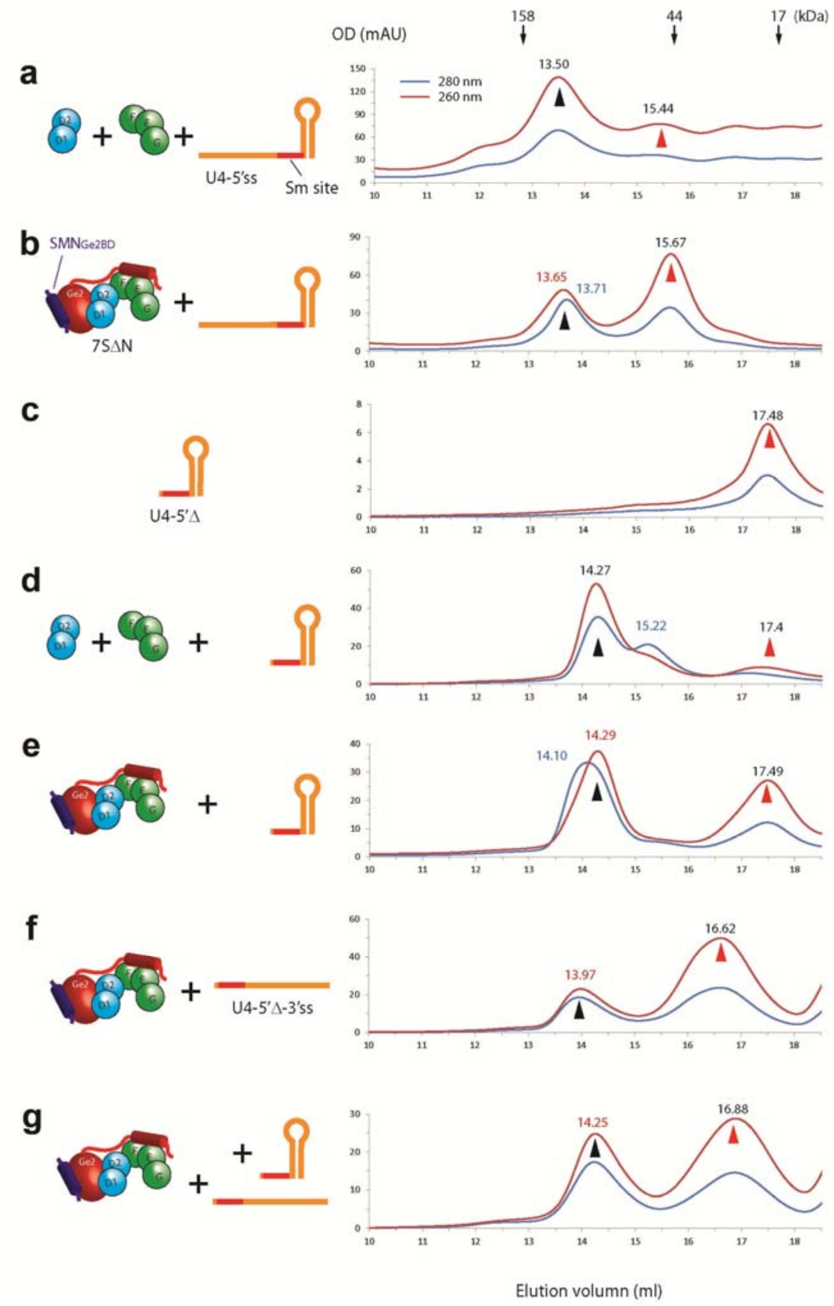

**Fig. S5. The gel filtration profiles of RNA and Sm subcore formation.** Mixture of 5Sm and U4-5'ss (a), mixture of 7SΔN and U4-5'ss (b), U4-5'Δ alone (c), mixture of 5Sm and U4-5'Δ (d), mixture of 7SΔN and U4-5'Δ (e), mixture of 7SΔN and U4-5'Δ-3'ss (f), or mixture of 7SΔN and equal molar amount of U4-5'Δ and U4-5'Δ-3'ss (g) was separated by gel filtration chromatography (right panels). For each, one representative result from at least two independent experiments is shown. (b) and (e) are the same as in Fig. 2 and are showed here for easy comparison. The input components are showed in cartoon (left panels). The peaks pointed at by the red arrow heads are the RNA peaks, and the peaks pointed at by the black arrow heads are the Sm-RNA complexes (subcores).

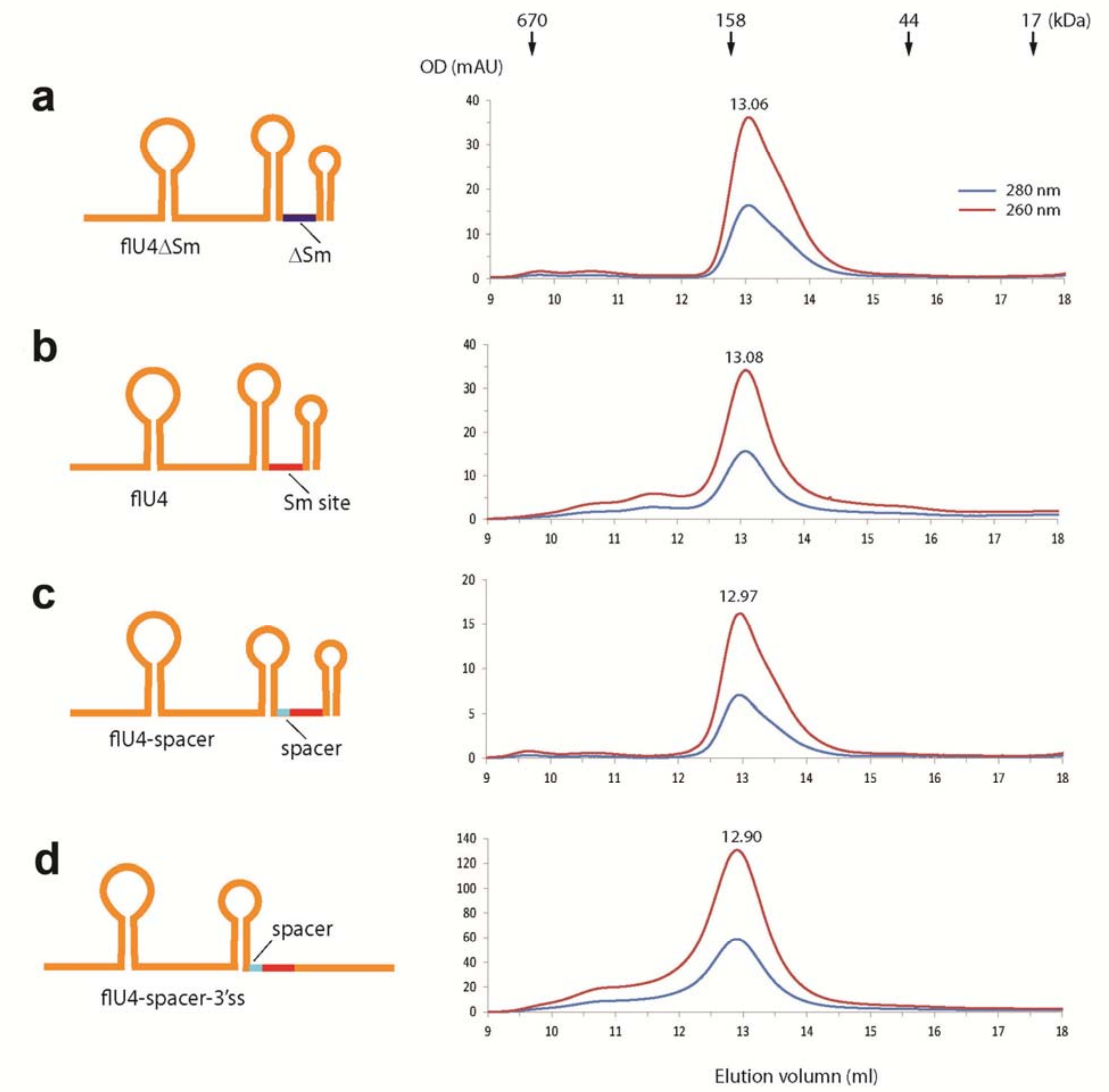

**Fig. S6. The gel filtration profiles of the four large-sized RNAs.** flU4ΔSm (a), flU4 (b), flU4-spacer (c) or flU4-spacer-3'ss (d) (in cartoon, left panel) was separated by gel filtration chromatography (right panels). For each, one representative result from two independent experiments is shown. The positions of standard proteins are indicated at the top.

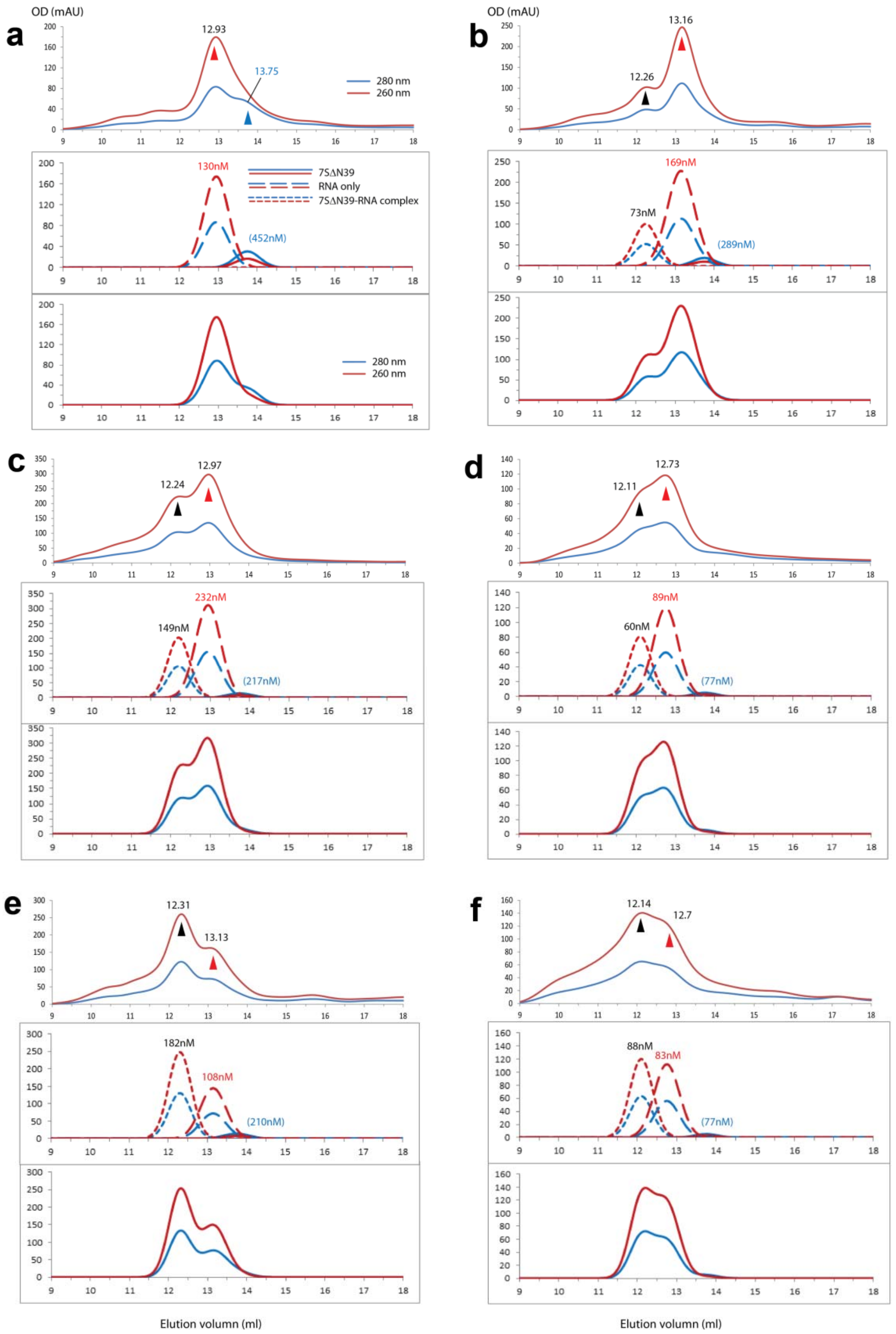

**Fig. S7. The simulations of GFC traces of component peaks and the combined traces in Fig 4.** Each panel corresponds to the same one in Fig.4. In each panel, the top traces are the experimental traces, the same as in Fig.4 (shown here for easy comparison), the middle traces are the simulated component peaks (and their concentrations, see Table S6 for the calculation.) and the bottom traces are the combined ones.

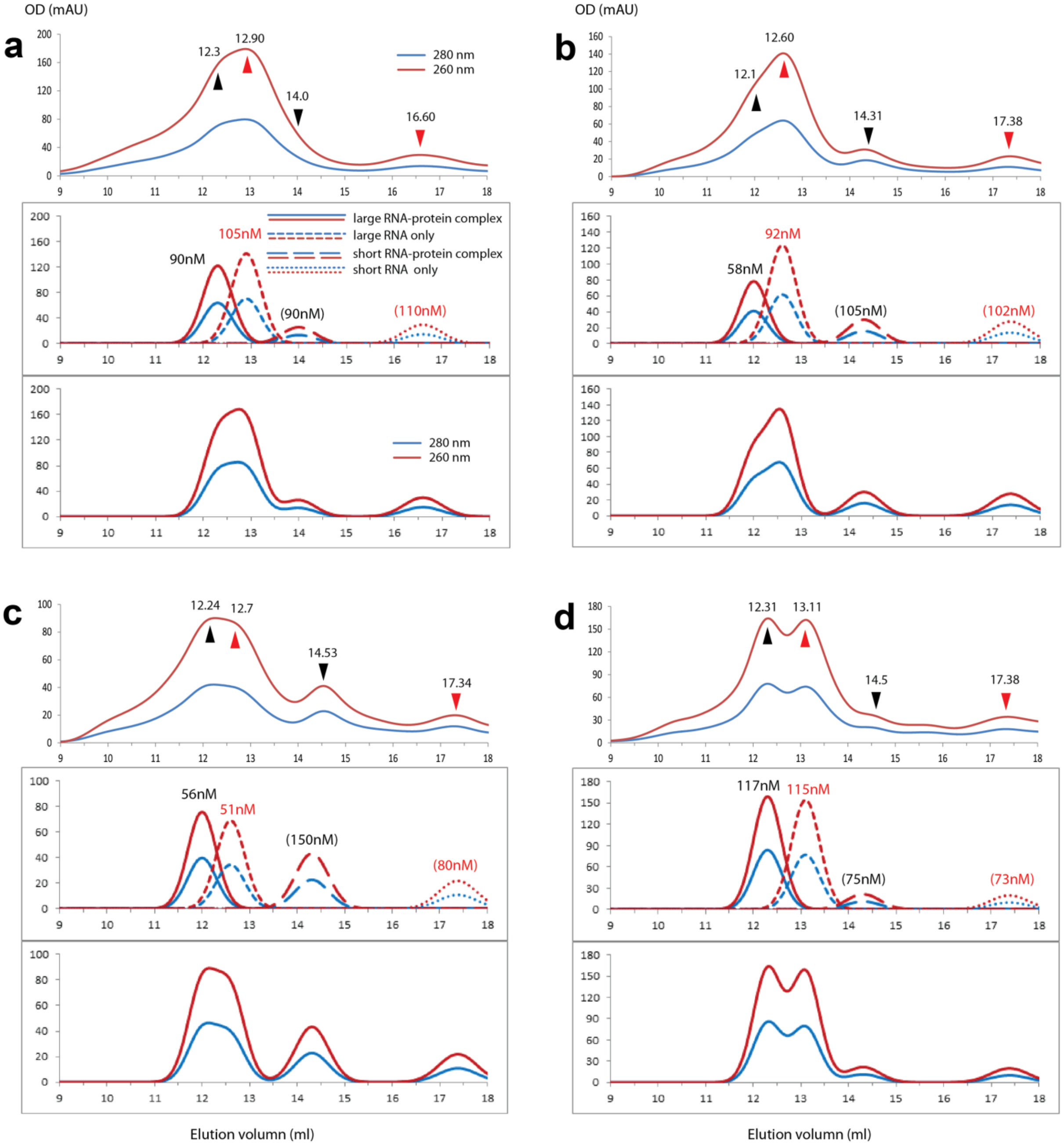

**Fig. S8. The simulations of GFC traces of component peaks and the combined traces in Fig 5.** Each panel corresponds to the same one in Fig.5. In each panel, the top traces are the experimental traces, the same as in Fig.4 (shown here for easy comparison), the middle traces are the simulated component peaks (and their concentrations, see Table S7 for the calculation.) and the bottom traces are the combined ones. Compared with the traces of the RNAs and RNA-protein complexes, the trace of 7SΔN is too small to contribute to the overall shape of the combined traces and is not shown in each panel.
